# Molecular mechanisms underlying the formation of larval green color and camouflage patterns in swallowtail butterfly, *Papilio memnon*

**DOI:** 10.1101/2023.05.18.541393

**Authors:** Liang Liu, Shinya Komata, Kai Wu, Tetsuya Kojima, Haruhiko Fujiwara

**Affiliations:** Department of Integrated Biosciences, Graduate School of Frontier Sciences, The University of Tokyo, Kashiwa, Chiba, 277-8562, Japan; School of Life Science and Technology, Tokyo Institute of Technology, Meguro-ku, Tokyo 152-8550, Japan; College of Life Sciences, Shangrao Normal University, Shangrao, China

**Author notes:** These authors contributed equally to this work and share first authorship (Liang Liu, Shinya Komata).

## Abstract

Insects have various strategies like mimicry or camouflage to avoid predation. Swallowtail butterfly larvae switch from a black and white pattern mimicking bird droppings to a green camouflage pattern in the fifth (final) instar. This larval pattern switch is regulated during the juvenile hormone (JH)-sensitive period, when JH titer declines rapidly, and *clawless* (*cll*), *abdominal-A* (*abd-A*), and *Abdominal-B* (*Abd-B*) function during this period. However, the molecular mechanism behind the background green color, a crucial aspect of the camouflage pattern, remains poorly understood. Here, we used *Papilio memnon*, which switches to the camouflage pattern in the fifth instar but is greenish from the third instar, to investigate the mechanism of camouflage pattern formation, particularly the larval green coloration.

Through RNA sequencing, we found that *BBP*s forming a gene cluster are upregulated in the green regions of *P. memnon* larvae during the fourth instar, whereas *P. xuthus* larvae, which have not yet turned green, showed minimal *BBP*s expression. When *BBP1* and *BBP2*, which were particularly highly expressed, were knocked down by RNAi, there was a phenotypic change in green to yellow in both fourth and fifth instar larvae. Expression analysis and knockdown experiments were conducted also for *JHBP*, which had been previously reported, and confirmed that it is involved in the synthesis of yellow pigment. Furthermore, knockdown of *Ubx* resulted in no phenotypic change in fourth instar larvae, but in fifth instar larvae, the eyespots pattern characteristic of the camouflage pattern almost entirely disappeared, suggesting that *Ubx* is also functional only during JH-sensitive period.

Our results indicate that the switch from mimetic to camouflage patterns resulted from the function of *cll*, *abd-A*, *Abd-B*, and *Ubx* prepatterning genes during the JH-sensitive period. And the increased expression of *BBP*s and *JHBP*s, independent of the JH-sensitive period, contributed to the development of green coloration.

## Introduction

Insects have a variety of adaptive traits to resist predation pressure. Butterflies are attractive prey for birds and other predators, but they often have a clever color pattern to avoid detection by predators through mimicry or camouflage (Pasteur, 1982; Futahashi & Fujiwara, 2008). Recent studies have highlighted the molecular and genetic mechanisms of wing color pattern formation involved in Müllerian or Batesian mimicry (Nijhout 1991; True et al. 1999; Joron et al. 2011; Kunte et al. 2014; Nishikawa et al. 2015; Mazo-Vargas et al. 2017; Matsuoka and Monteiro 2018). The color patterns of *Papilio* butterfly larvae, such as *Papilio xuthus*, also show some intriguing body patterns. The larvae of *Papilio* butterflies are known to dramatically change their color pattern from a bird-dropping pattern (mimetic pattern) to a camouflage pattern (Prudic et al. 2007; Futahashi and Fujiwara 2008). The first to the fourth instar larvae show black and brown color patterns with white V-shaped patches (mimetic pattern), but the fifth (last) instar larvae have a greenish body with a pair of false eyespots on the third thoracic segment (T3) and a V-shaped marking on the abdominal segments (camouflage pattern). The dramatic color pattern change from the fourth to the fifth instar larvae has been the subject of many molecular mechanism studies, and it has been reported that a rapid decrease in juvenile hormone (JH) titer at the beginning of the fourth instar larvae, i.e., immediately after the third molt, causes a switch from mimetic pattern up to the fourth instar larvae to the fifth instar larvae’s camouflage pattern (Futahashi and Fujiwara 2008). The period up to about 12 hours after the third molt, when JH titer decreases, is called the JH sensitive period (JHSP) (Futahashi and Fujiwara 2008; Jin et al. 2019).

The larval color pattern switch is hypothesized to comprise two independent processes: the prepatterning process during the JHSP and the pigmentation process during the molting period (Jin et al. 2019). The prepatterning process is speculated to define the coarse boundary of the segment-specific pattern like drawing some dashed lines on the larval body, and the synthesis or transportation of the downstream pigments will be initiated in the predefined region during the pigmentation process like painting. Recently, a more specific JH-dependent network involving three homeobox genes (i.e., *clawless* (*cll*), *abdominal-A* (*abd-A*), and *Abdominal-B* (*Abd-B*)) was reported to control the prepatterning process during the JHSP in *P. xuthus* (Jin et al. 2019). *cll* is involved in the formation of the eyespot pattern on T3 of fifth instar larvae, and *abd-A* and *Abd-B* are involved in the formation of the V-shaped marking on the abdominal segments (Jin et al. 2019). On the other hand, there are many known pigmentation genes that would be involved in the pigmentation process. *Tyrosine hydroxylase* (*TH*), *dopa decarboxylase* (*DDC*), *yellow*, *tan*, and *laccase 2* are associated with the larval cuticular black coloration, but another melanin-related gene *ebony* which converts dopamine into N-beta alanyl dopamine (NBAD), is responsible for the red coloration in the eyespots of *P. Xuthus* (Futahashi and Fujiwara 2005; Futahashi and Fujiwara 2006; Futahashi and Fujiwara 2007; Futahashi et al. 2010; Shirataki et al. 2010; Futahashi et al. 2012). Not only the red and black pigments (melanin) in eyespots of the third thoracic segment, the *Papilio* larval body also shows a background green color, which serves as cryptic camouflage to hide the caterpillars in the host plants. In *P. xuthus*, the expression of pigmentation genes associated with green is thought to be initiated in JHSP, and it has been hypothesized that JH titer regulates the pigmentation gene expression (Futahashi and Fujiwara 2008; Jin et al. 2019), but the relationship between pigmentation genes and JH titer is not fully understood. The commonly observed larval green coloration in Lepidopteran species usually results from a combination of bilin pigment (blue) and carotenoids (yellow) (Riley et al. 1984; Shirataki et al. 2010; Futahashi et al. 2012). The expression of *bilin-binding protein*s (*BBP*s) and *carotenoid-binding protein 1* (*CBP1*) or yellow-related genes in *P. xuthus* and *P. polytes* showed high consistency with the distribution of the larval blue and yellow coloration, respectively (Futahashi et al. 2012). Additionally, a previous study suggested that larval and pupal protective green color of *P. polytes* resulted from the functions of *BBP*s contributing to the blue coloration and *juvenile hormone-binding protein*s (*JHBP*s) responsible for the yellow coloration (Yoda et al. 2020).

Both *BBP*s and *JHBP*s form gene clusters with 6 to 8 genes in close proximity on the same chromosome; in *P. polytes* larvae and pupae, only some genes in clusters of *BBP*s and *JHBP*s are involved in blue and yellow formation (Yoda et al. 2020). On the other hand, in the closely related species *P. memnon*, the color pattern switch from 4th to 5th instar larvae occurs as in *P. xuthus* and *P. polytes*, but 3rd and 4th instar larvae already have greenish body color instead of black or brown (fig. 1A). Comparing the body coloration of fourth instar larvae of *P. xuthus* and *P. memnon*, *P. memnon* is clearly greenish (fig. 1A). Although *P. memnon* has a larger body size compared to *P. xuthus* and *P. polytes* (*P. memnon* pupae: 1.5–3.5 g, *P. xuthus* pupae: 0.5–1.5 g (Komata and Sota 2017; Komata et al. 2018) differences in habitat and behavior, and other differences are not clear, and the adaptive significance of *P. memnon*’s 3rd and 4th instar larvae already being greenish is not known. In addition, the molecular mechanisms involved in the color pattern formation of *P. memnon* larvae have not been studied at all. In this study, we investigated the molecular mechanisms of color pattern formation in *P. memnon* larvae and compared them with those of *P. xuthus* and *P. polytes* in order to provide insight into the elucidation of the adaptation and evolutionary mechanisms of color patterns in larvae of *Papilio* butterflies. First, JH-analog application experiments were conducted to investigate whether the reduction of JH titer, as in *P. xuthus* (Futahashi and Fujiwara 2008), is responsible for the color pattern switch from 4th to 5th instar larvae in *P. memnon*. Next, we searched for genes associated with larval green coloration by RNA sequencing (RNA-seq) and compared the expression levels of pigmentation genes such as *BBP*s and *JHBP*s in fourth instar larvae of *P. xuthus* and *P. memnon*. In addition, RNAi experiments for *BBP*s and *JHBP*s were performed by *in vivo* electroporation-mediated method (Ando and Fujiwara 2013) to determine which *BBP*s and *JHBP*s in the clusters are involved in the formation of larval green color in *P. memnon*. We also performed RNAi experiments for *Ultrabithorax* (*Ubx*) because RNA-seq data showed that the homeobox gene, *Ubx*, is highly expressed in the anterior region of larvae in *P. memnon*. The molecular mechanisms and adaptive significance of the greenish body coloration of *P. memnon* larvae from the 3rd and 4th instar larvae are discussed in comparison with *P. xuthus* and *P. polytes*.

**Figure 1.**
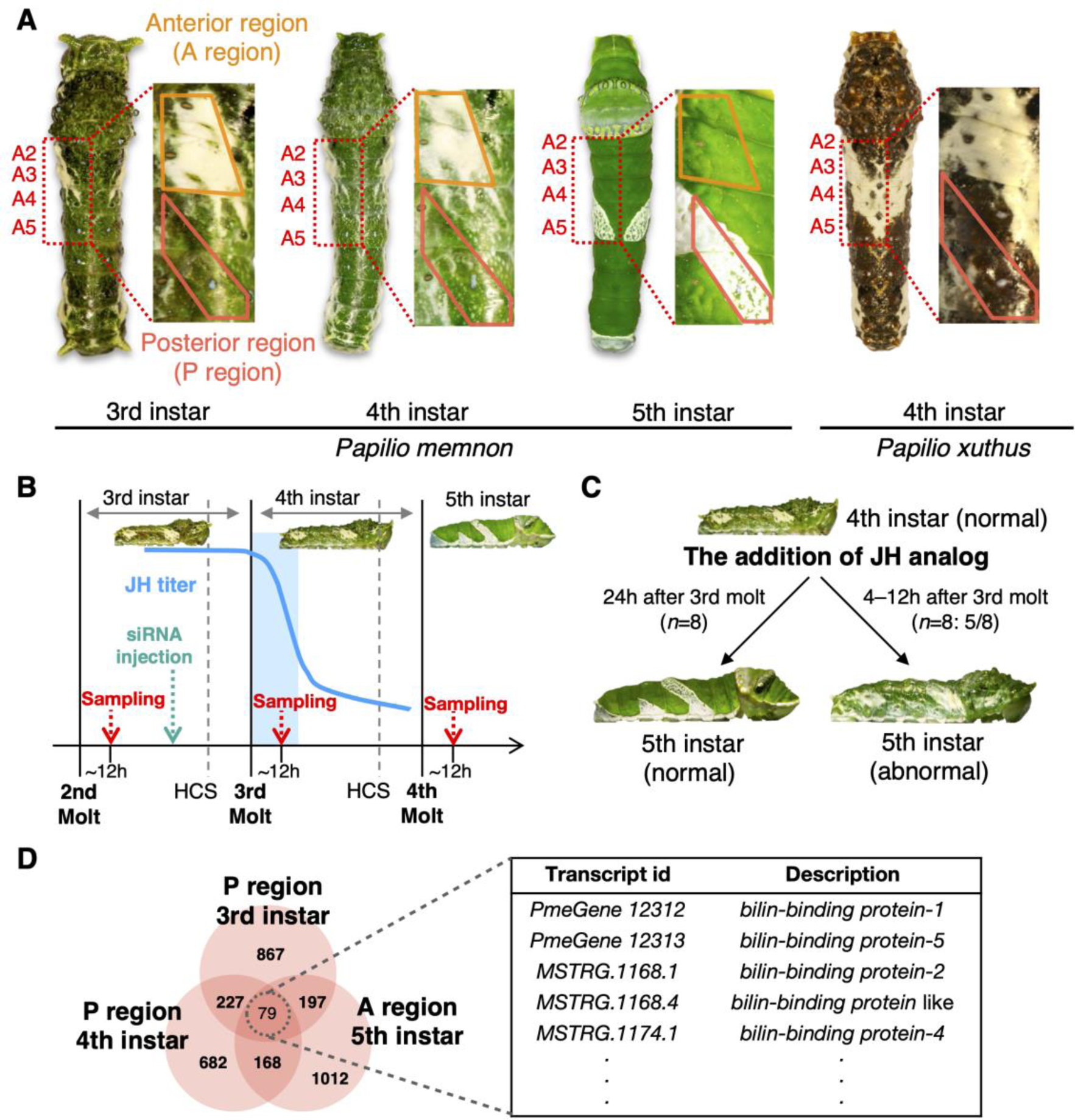
Larval color patterns and underlying juvenile hormone (JH) titers, and the search for green color-associated genes by RNA sequencing (RNA-seq) in *Papilio memnon*. (A) Dorsal and magnified lateral view of third, fourth and fifth instar larvae of *P. memnon* and fourth instar larvae of *P. xuthus* are shown in the left and right panel, respectively. The orange and pink frames represent two regions of epidermal tissues collected for RNA-seq. The anterior region (A region, across the second and third abdominal segment) shows white color in the third and fourth instar, but switches to green color in the fifth instar, The posterior region (P region, across the fourth and fifth abdominal segment) shows in green color in the third and fourth instar, but switch to white color in fifth instar. (B) The expected JH titer in *Papilio* butterflies are shown by blue curve (Futahashi and Fujiwara 2008). The blue region indicates the JH-sensitive period (JHSP, the first 12 h after the 3rd molt) during which JH titer drops rapidly (Futahashi and Fujiwara 2008). HCS: head capsule slippage. We performed RNA sampling for RNA-seq at the timing indicated by the red arrow (12 h after molts) and siRNA injection for RNAi at the timing indicated by the green arrow. (C) Similar to the previous study on *P. xuthus* (Futahashi and Fujiwara 2008), when JH-analog was applied to larvae during JHSP, even fifth instar larvae in *P. memnon* showed the same color pattern as fourth instar larvae. (D) RNA-seq revealed 79 genes highly expressed in the green region of 3rd to 5th instar larvae (P region in 3rd and 4th instar larvae and A region in 5th instar larvae) in *P. memnon*. 79 genes were searched for genes involved in pigment synthesis, and several *bilin binding protein* (*BBP*) genes involved in blue pigment synthesis were found.

## Results

### JH-analog application and green related gene screening by RNA-seq

First, we applied JH-analog to *P. memnon* larvae to confirm whether the switch from a mimetic pattern (up to 4th instar larvae) to a camouflage pattern (5th instar larvae) occurs due to a decrease in JH titer immediately after the third molt (fig. 1B), as in the case of *P. xuthus* (Futahashi and Fujiwara 2008). All larvae that were exposed to JH-analog 24 hours after the third molt switched normally to the camouflage pattern (*n*=8). But when JH-analog was applied 4 and 12 hours after the third molt, most 5 out of 8 larvae had the mimetic pattern despite being 5th instar larvae (fig. 1C and supplementary fig. S1). This result is similar to the JH-analog application experiment in *P. xuthus* (Futahashi and Fujiwara 2008), indicating that the decrease in JH titer between 12 and 24 hours after the third molt is responsible for the switch from the mimetic pattern to the camouflage pattern in *P. memnon*. Next, to search for genes involved in the green coloration of *P. memnon* larvae, we searched for genes highly expressed in the green region of larvae by RNA-seq (Supplementary table S1). First, we extracted the different expressed genes (DEGs) highly expressed in the green region (posterior region of 3rd and 4th instar larvae, anterior region of 5th instar larvae, fig. 1A) for each instar, and then found 79 genes that were commonly highly expressed in the green region of all three instars (fig. 1D; Supplementary Table S2. Among the 79 genes, we identified 11 cuticle proteins and piment-related genes and found that they contain 5 *BBP* genes (fig. 1D). To be more specific, 29 basis cellular events-associated genes, 9 signaling pathway-related genes, 4 transcriptional factors, 6 cuticular protein-related genes, and 5 pigment-binding protein-related genes were identified. Among the 29 basis cellular events-associated genes, for example, *MSTRG.10015.1* was characterized as *filaggrin-2-like*, which is essential for normal cell-cell adhesion in the cornified cell layers. In the 9 signaling pathway-related genes, *discoidin domain-containing receptor 2-like* (*PmemnonGene0006583.mrna1*) functions as cell surface receptor for fibrillar collagen and regulates cell differentiation. And *PmemnonGene0000673.mrna1*, which was identified as *sorting nexin lst-4*, is involved in the signaling of vulval development by acting as a negative regulator of epidermal growth factor receptor (EGFR) signaling. All the 5 pigment-binding protein-related genes identified in the 79 green color-associated DEGs appeared to be Bilin-binding proteins which are involved in the insect blue coloration.

### Clusters and expression analysis of genes associated with green color

We found that *BBP*s highly expressed in the green region are genes in a gene cluster of *BBP*s that have already been reported in other *Papilio* butterflies (Yoda et al., 2020; fig. 2A). The gene cluster of *BBP*s was found to have six *BBP*s in *P. memnon*, in the same arrangement and orientation as in *P. polytes*, *P. xuthus* and *P. machaon* (fig. 2A and supplementary fig. S2). Furthermore, when the expression levels of *BBP*s in 3rd, 4th, and 5th instar larvae were examined based on RNA-seq data, the expression of *BBP1* and *BBP2* tended to be particularly high in the green region (anterior region) of 5th instar larvae (fig. 2B). The expression of *BBP4*, *BBP5*, and *BBP6* was higher in the green region (posterior region) of 4th instar larvae than in other stages and regions, although the overall expression level was not large.

**Figure 2.**
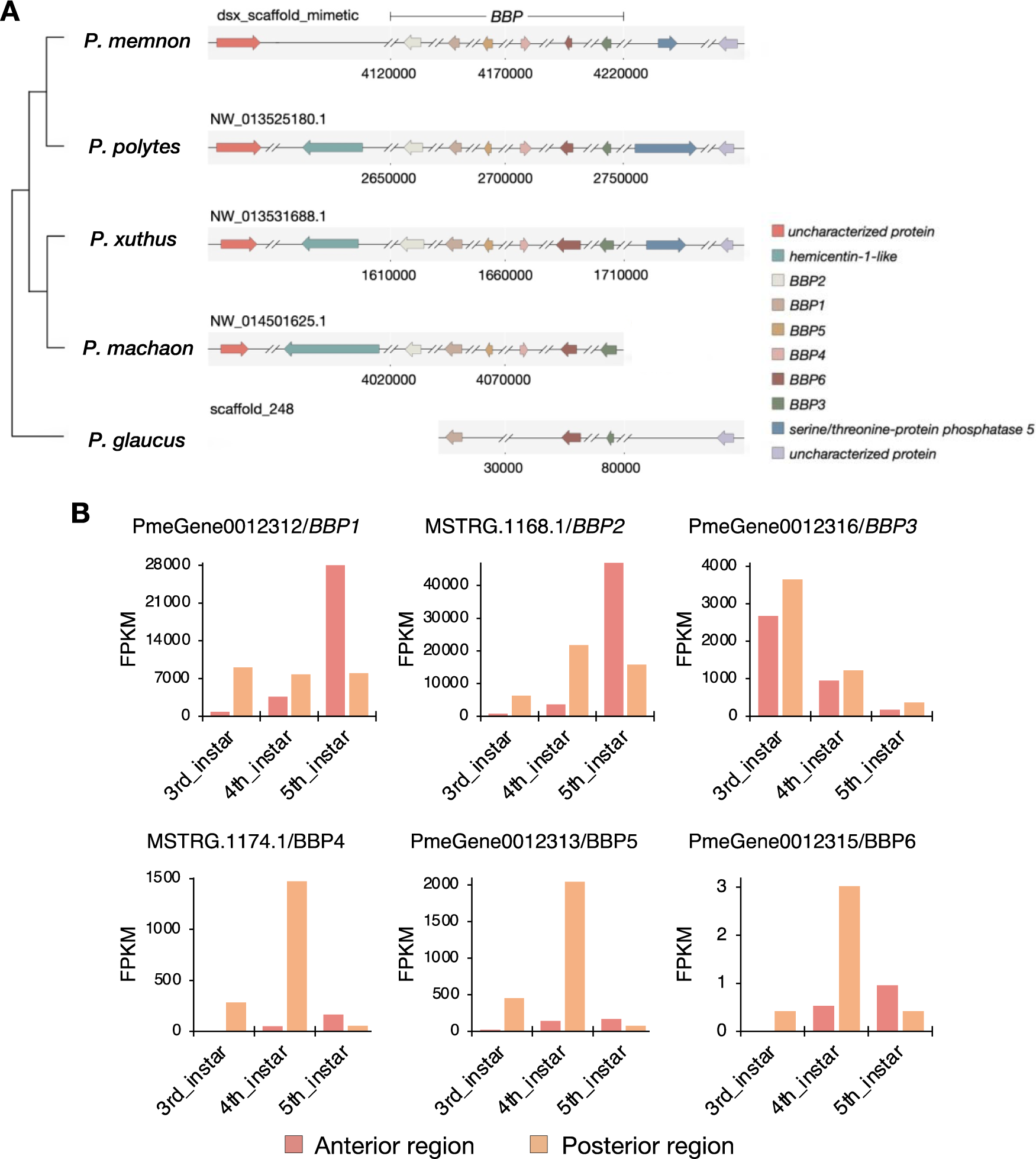
Structure and expression profiles of *bilin-binding protein*s (*BBP*s) (A) Cluster of *BBP*s in five *Papilio* butterflies. The length of arrows represents the relative gene length. Transcriptional orientation is shown by the direction of arrows. Gene clusters with six *BBP*s were identified in all four species except *P. glaucus*. (B) Expression profiles of *BBP*s in anterior and posterior regions (fig. 1A) of *P. memnon* by RNA-seq from 3rd to 5th instars (*n* =1 but one sample is mixed with three individuals). The expression level of *BBP*s is shown by the fragments per kilobase of exon per million mapped fragments (FPKM).

Next, we compared the expression of *BBP*s in the posterior region of 4th instar larvae (green region in *P. memnon* and black/brown region in *P. xuthus*, fig. 1A) between *P. memnon* and *P. xuthus*, and found that all six *BBP*s were highly expressed in *P. memnon*, but almost none in *P. xuthus* (fig. 3). In addition, the expression of *TH*, *DDC*, *tan*, *yellow*, and *Laccase2*, which are involved in melanin synthesis, was also compared between *P. memnon* and *P. xuthus*, and these genes were found to be expressed without significant differences between the two species (fig. 3).

**Figure 3.**
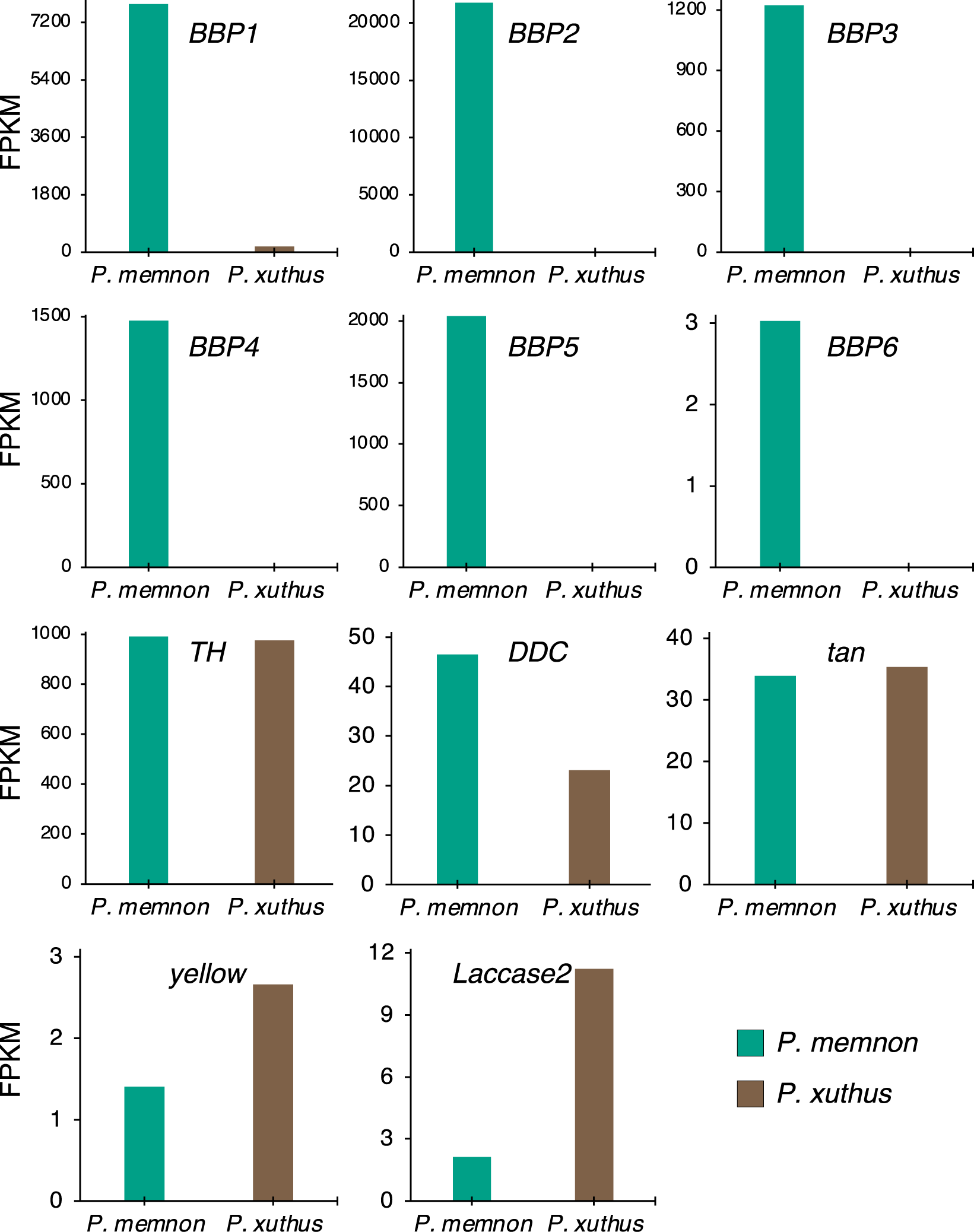
Comparison of expression levels of *bilin binding protein*s (*BBP*s) and melanin-related genes in *Papilio memnon* and *Papilio xuthus* by RNA-seq data in the posterior regions of 4th instar larvae. The expressions of all six *BBP*s were significantly higher in *P. memnon* than *P. xuthus*, but those of melanin-related genes (i.e., *TH*, *DDC*, *tan*, *yellow*, *Laccase2*) were not significantly different. The expression level is shown by the fragments per kilobase of exon per million mapped fragments (FPKM).

Furthermore, we examined the expression and clustering of *JHBP*s in *P. memnon*, which is involved in the synthesis of yellow pigment in *P. polytes* and was reported to have a gene cluster similar to that of *BBP*s. We found that eight *JHBP*s also formed a gene cluster in *P. memnon*, and that *JHBP4812*, *JHBP4813*, and *JHBP4815* were highly expressed in the anterior region (green region) of fifth instar larvae (supplementary figs. S2 and S3), and compared to *P. xuthus*, *JHBP4811*, *JHBP4812*, *JHBP48813* and *JHBP4815* were found to be highly expressed in *P. memnon* (supplementary fig. S3).

### Functional analysis of *BBP*s and *JHBP*s in the larval green region

*BBP1* and *BBP2*, which were particularly highly expressed in the *BBP* gene cluster, were knocked down by electroporation-mediated RNAi (Ando and Fujiwara 2013). siRNA of *BBP1* and *BBP2* was introduced in third instar larvae, and a clear change in the green region to yellow was observed (figs. 4A and B and supplementary figs. S4 and S5). The yellowish change indicates that the synthesis of the blue pigment was blocked. Interestingly, a similar green-to-yellow change was observed not only in fifth instar larvae but also in fourth instar larvae (figs. 4A and B and supplementary figs. S4 and S5). When siRNAs of *BBP3* and *BBP5* were injected at the same time (double knockdown), the green region turned slightly yellowish (fig. 4C and supplementary fig. S6). However. the knockdown of *BBP4* and *BBP6* didn’t cause any change on the larval green background color (fig. S7). Previous study showed that *cll*, *abd-A*, and *Abd-B*, which play important roles in switching from mimicry to camouflage patterns, were knocked down by siRNA injection in 3rd instar larvae, but no phenotypic change was observed in 4th instar larvae, and phenotypic change was observed only in 5th instar larvae (Futahashi and Fujiwara 2008; Jin et al. 2019). This means that these prepatterning genes are important only for JH-sensitive period, but for *BBP1* and *BBP2*, they are expressed and involved in the synthesis of larval blue pigment regardless of JH-sensitive period.

**Figure 4.**
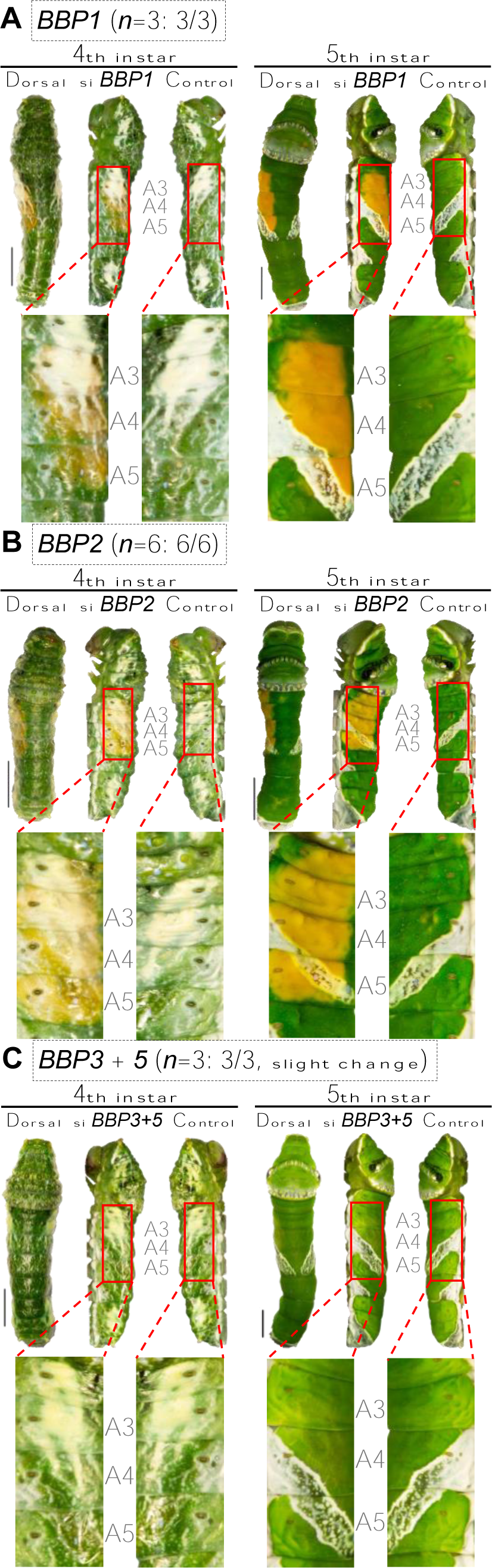
*in vivo* electroporation-mediated RNAi of *BBP*s. (A) 4th-instar (left) and 5th-instar (right) larvae after RNAi of *BBP1*. (B) 4th-instar (left) and 5th-instar (right) larvae after RNAi of *BBP2*. (C) 4th-instar (left) and 5th-instar (right) larvae after multiple-RNAi of *BBP3* and *BBP5*. siRNA is injected through the intersegmental membrane between the 7th and 8th abdominal segment and introduced into specific epidermal region (A3–5) via an electroporation-mediated method during the 3rd instar. scale bar = 5 mm. Knockdown of *BBP1* and *BBP2* caused complete elimination of blue color (A, B) while the double knockdown of *BBP3* and *BBP5* only slightly reduce the blue color(C). The number of samples showing phenotypic changes among the tested numbers is shown in parentheses (e.g., 3/3 for *BBP1* knockdown). Supplementary figures S4 and S5 show other replicates.

Knockdown of *JHBP4815* changed the green region to bluish, possibly blocking the synthesis of yellow pigment, and phenotypic changes were observed in both 4th and 5th instar larvae as those in *BBPs* (fig. 5A and supplementary fig. S8). Although no clear phenotypic changes were observed in larvae by the other *JHBP*s knockdowns (fig. 5B and supplementary fig. S9), knockdowns of *JHBP4811* and *JHBP4813* caused pupal green to turn slightly blue, suggesting that yellow synthesis was inhibited in *P. memnon* pupae (supplementary fig. S10). *JHBP4811* and *JHBP4813* were reported to be involved in pupal green formation (synthesis of yellow pigment) in *P. polytes* (Yoda et al. 2020). For all of the above knockdown experiments, we confirmed reduced expression of the target genes with statistical significance in the knocked-down regions by RT-qPCR as described in Jin et al., 2019 (supplementary fig. S11, all primers used in this study are listed in Supplementary table S4).

**Figure 5.**
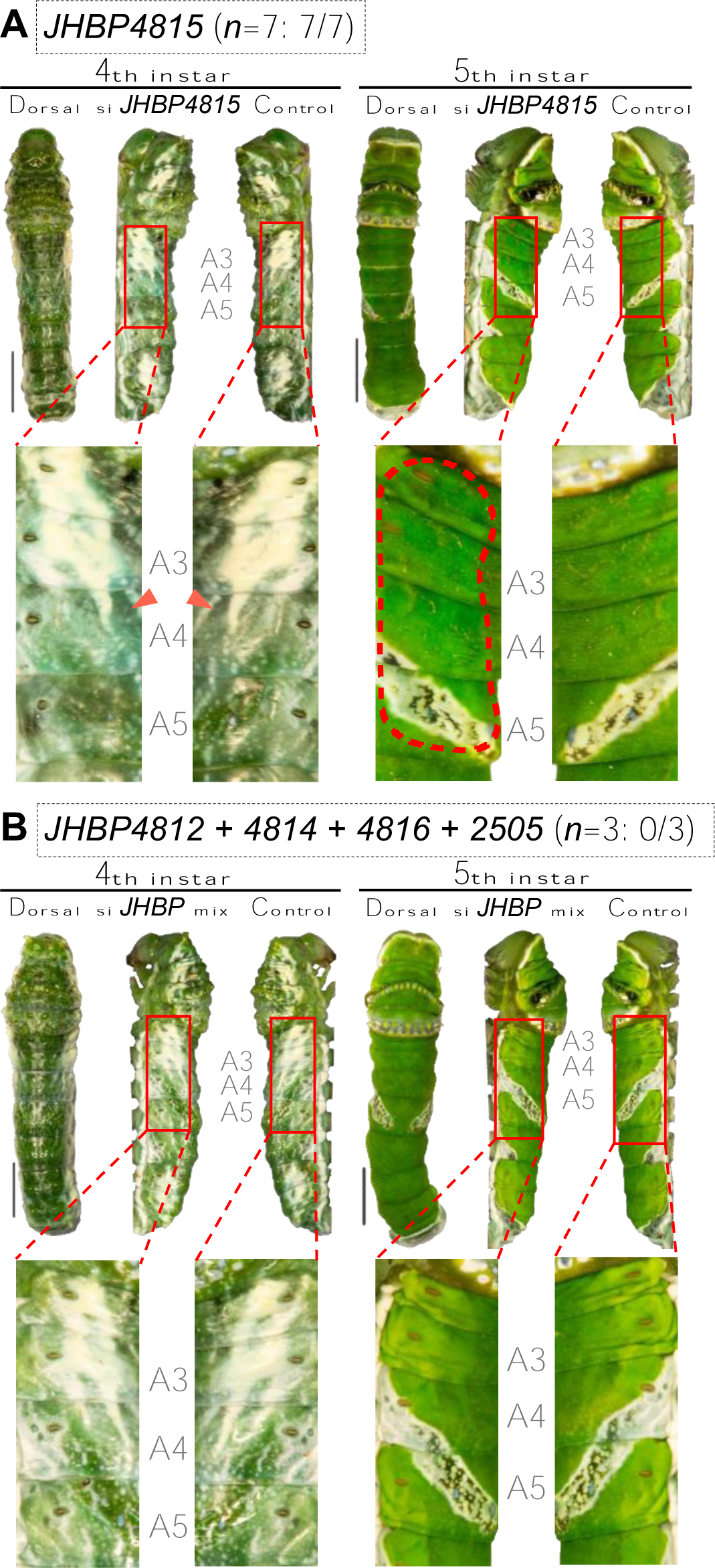
*in vivo* electroporation-mediated RNAi of *JHBP*s. (A) 4th-instar (left) and 5th-instar (right) larvae after RNAi of *JHBP4815*. (B) 4th-instar (left) and 5th-instar (right) larvae after multiple-RNAi of *JHBP4812*, *JHBP4814*, *JHBP4816* and *JHBP2505*. siRNA is injected through the intersegmental membrane between the 7th and 8th abdominal segment and introduced into specific epidermal region (A3–5) via an electroporation-mediated method during the 3rd instar. scale bar = 5 mm. Among all the RNAi experiments targeting *JHBP*s, only knockdown of *JHBP4815* caused a slight reduction of yellow color during the larval stage. The number of samples showing phenotypic changes among the tested numbers is shown in parentheses (e.g., 7/7 for si *JHBP4815*). Supplementary figure S8 shows other replicates.

### Newly discovered prepatterning gene *Ubx* in *P. memnon* larvae

*cll*, *abd-A*, *Abd-B* and *delta* have been reported in *P. xuthus* as prepatterning genes involved in the switch from mimicry to camouflage patterns (Jin et al. 2019; Jin et al. 2020). We first identified the orthologs of these prepattening genes in *P. memnon* (fig. S12) and then tested whether they have similar functions in *P. memnon* was confirmed by expression analysis using RT-qPCR and gene knockdown using RNAi (supplementary figs. S13 and S14). We confirmed that *cll* is involved in the formation of the T3 eyespots pattern and *abd-A Abd-B* and *delta* are involved in the formation of the posterior V-shaped pattern, as in *P. xuthus* (supplementary fig. S14). However, the phenotypic change in the eyespots pattern of T3 due to knockdown of *cll* is not so large in our results in *P. memnon* and in the results of *P. xuthus* in the previous study: only a part of the eyespots pattern is lost (supplementary fig. S14; Jin et al., 2019). Therefore, it is conceivable that there may be other prepatterning genes responsible for the eyespot formation. Then we knocked down *Ubx* in *P. memnon* larvae, since RNA-seq analysis showed that the homeobox gene *Ubx* was highly expressed in the anterior region (supplementary fig. S15) and RT-qPCR also showed that its expression was increased in the anterior region of larvae (supplementary fig. S13). As a result, when siRNA of *Ubx* was introduced in 3rd instar larvae, no phenotypic change was observed in 4th instar larvae, but in 5th instar larvae, almost all of the eyespot pattern in T3 disappeared (fig. 6A and supplementary fig. S16). Furthermore, when *Ubx* siRNA was also introduced in the first abdominal segment (A1) next to T3, no phenotypic change was observed in 4th instar larvae, but almost all ramen-like patterns disappeared in 5th instar larvae (fig. 6B and supplementary fig. S17). These results indicate that *Ubx* functions in JH-sensitive period and is an important prepatterning gene for the switch from mimetic to camouflage patterns.

**Figure 6.**
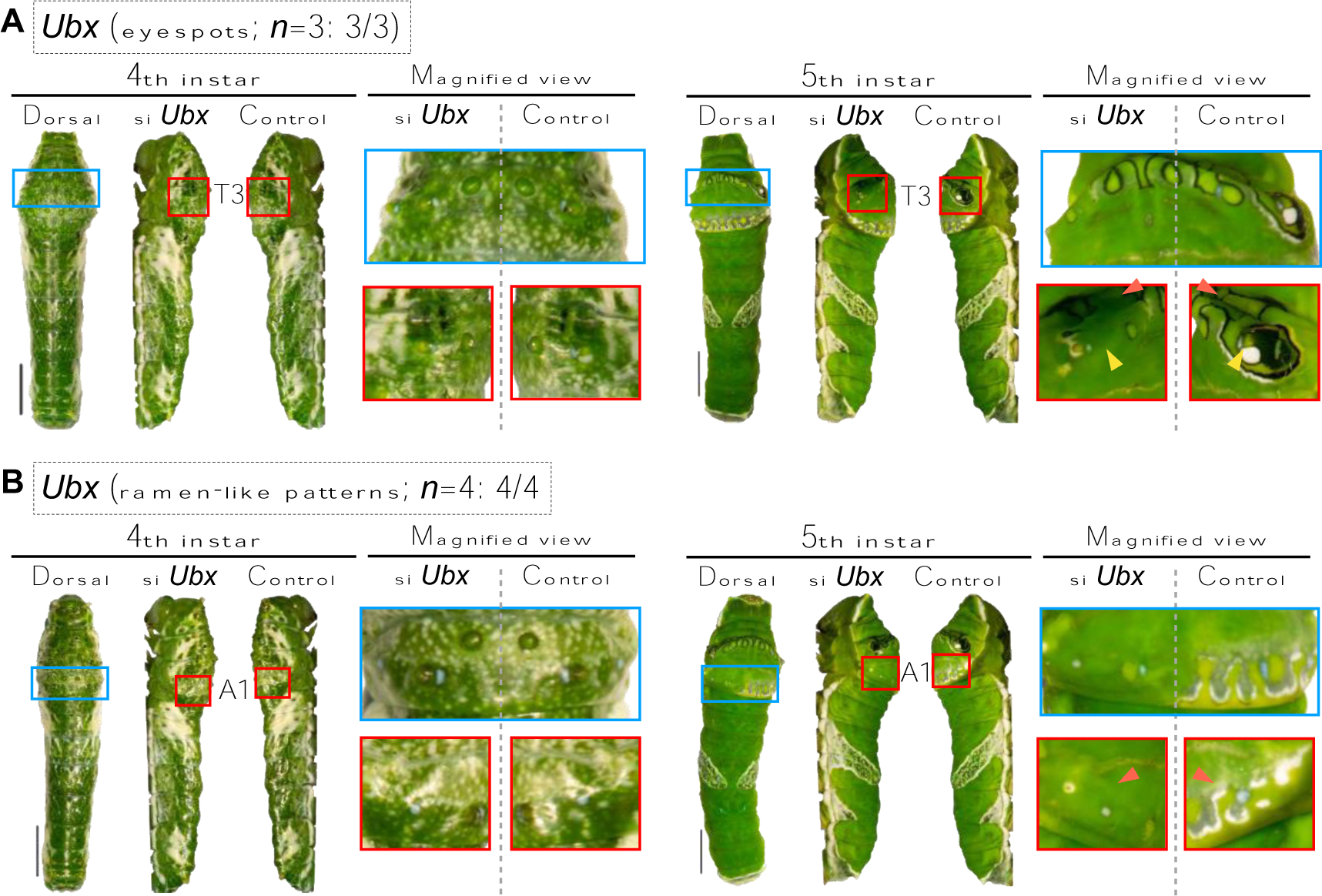
*in vivo* electroporation-mediated RNAi of *Ubx*. (A) *Ubx* knockdown on the third thoracic segment (T3). (B) *Ubx* knockdown on the first abdominal segment (A1). siRNA is injected through the intersegmental membrane between the 7th and 8th abdominal segment, and introduced into specific epidermal regions via an electroporation-mediated method during the 3rd instar. scale bar = 5 mm. Eyespots on segment T3 and ramen-like patterns on segment A1 were completely removed by the knockdown of *Ubx*. Supplementary figures S16 and S17 show other replicates.

## Discussion

In the present study, it was found that *BBP*s and *JHBP*s turn larvae green (fig. 7), and it is thought that the expression of *BBP*s and *JHBP*s begins earlier in *P. memnon* than in *P. xuthus* and *P. polytes*, so that *P. memnon* larvae turn green from the third and fourth instars (fig. 3 and supplementary fig. S3 C). I In *P. xuthus*, the switch to camouflage patterns and from black/brown to green occurs simultaneously, leading to the hypothesis that the expression of pigment synthesis genes such as *BBP*s and *JHBP*s begins during the JH-sensitive period (Futahashi and Fujiwata 2008; Jin et al. 2019). In *P. memnon*, on the other hand, there may be no relationship between the decrease in JH titer and the expression of *BBP*s and *JHBP*s, and some other factor may control the expression of these greenish-related genes. In addition, *BBP1* and *BBP2*, which are involved in the synthesis of blue pigment in both larvae and pupae of *P. polytes*, were found to have similar functions in *P. memnon* larvae in this study (fig. 7). *P. polytes* and other *Papilio* species have green, brown and orange polymorphisms in their pupae (Yoda et al. 2020), then the relationship between the mechanisms controlling pupal coloration and those controlling larval coloration is also interesting: even in the pupae of *P. polytes*, there is a significant difference in *BBP*s expression between green and brown pupae (Yoda et al. 2020). In other words, *BBP*s expressions are greater in green pupae. The pupal brown/green color switch is known to be related to pupal cuticle melanizing hormone (PCMH) (Awiti and Hidaka 1982; Starnecker and Hazel 1999; Yamanaka et al. 1999) and PCMH may upregulate *BBP*s and *JHBP*s expressions, but this is not well understood. Further studies are needed to investigate the underlying mechanisms that trigger the formation of green coloration, which is important for larval and pupal camouflage, i.e., the expression of *BBP*s and *JHBP*s.

**Figure 7.**
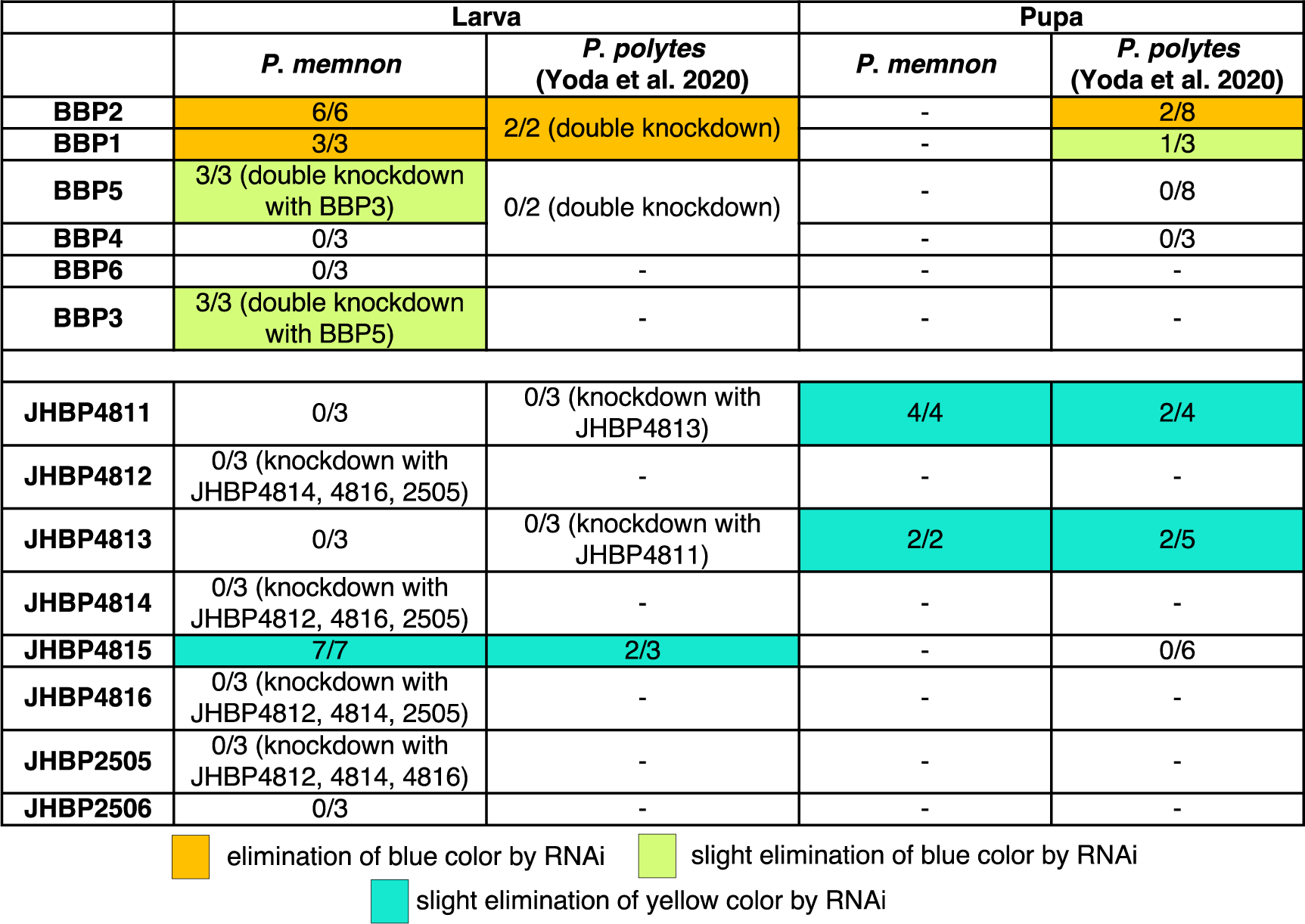
Summary of the results of RNAi experiments in *P. memenon* and *P. polytes* in this and previous studies (Yoda et al., 2020) *BBP1* and *BBP2*, whose phenotypes changed significantly by knockdown, are thought to be involved in larval green formation both in *P. memnon* and *P. polytes*. However, no phenotypic change in *BBP5*, which is slightly altered by knockdown in *P. memnon*, was observed in *P. polytes* (Yoda et al., 2020). *JHBP4815* is thought to have a common role in larvae and pupae of *P. memnon* and *P. polytes*.

For the knockdown of *JHBP*s, the phenotypic changes were not as significant as those seen with the knockdown of *BBP*s, with only a slight loss of yellow pigment. As for the yellow pigment, it is shown to be associated with the *carotenoid-binding protein 1* (*CBP1*) or yellow-related genes in *P. xuthus* and *P. polytes* (Futahashi et al. 2012; Yoda et al. 2020), and some putative carotenoid-binding proteins are characterized as members of the takeout/JHBP family. Therefore, it is not surprising that *JHBP*s are also involved in yellow pigment synthesis in *P. memnon*. Perhaps there are other important *JHBP*s or *CBP*s regulating the yellow pigment synthesis than the ones we have focused on in the *P. memnon* genome. Furthermore, there may be functional redundancy in *BBP*s and *JHBP*s, since even *BBP5* and *BBP3* showed small phenotypic changes due to knockdown.

Despite the fact that *BBP*s are speculated to be functionally redundant, in this study, RNAi of *BBP1* and *BBP2* both completely transformed the larval integument from green to yellow, suggesting the complete loss of blue pigments (fig. 4). A previous study suggested that the epidermal biliproteins formed a functional dimer in *Rhodinia fugax* (Saito 1998; Saito 2001). Thus, a possible explanation for the complete loss of blue color caused by the knockdown of *BBP*1 or *BBP*2 is that *BBP1* and *BBP2* also form a functional dimer in the larval epidermis of *P. memnon*, like *R. fugax*.

In addition, *JHBP*s differs from *BBP*s in that the genes involved in the synthesis of yellow pigment differ between larvae and pupae. That is, *JHBP4815* is involved in yellow pigment synthesis in larvae and *JHBP4811* and *JHBP4812* in pupae (figs. 5 and 7 and supplementary fig. S10). This finding suggests that multiple *JHBP*s are possible to be responsible for the yellow coloration in differential stages. Furthermore, *BBP3* and *BBP5* have not been shown to be involved in green formation in *P. polytes* (Yoda et al. 2020) but are involved in green formation in *P. memnon* (figs. 4 and 7). Such differences in the role of *BBP*s among species may also lead to differences in whether third- or fourth instar larvae are greenish or not. A more comprehensive comparative analysis of expression and function among species will be necessary in the future.

In this study, we confirmed a conserved JH-dependent prepatterning network involving three homeobox genes (*cll*, *abd-A* and *Abd-B*) and newly identified a novel prepatterning gene *Ubx* which is essential to the anterior pattern (eyespots and ramen-like pattern) formation in *P. memnon*. Intriguingly, the three prepatterning genes *Ubx*, *abd-A* and *Abd-B* belong to a homeotic complex called Bithorax (BX-C), primarily controlling the segmental identity and differentiation of the third thoracic segment and the first to the eighth abdominal segments (Karch et al. 1985). Null mutation of the homeotic gene *Ubx* caused a remarkable morphological change transforming the halteres into a pair of wings in *D. melanogaster* (Bender et al. 1983; Weatherbee et al. 1998; Hersh et al. 2007; Pavlopoulos and Akam 2011). The four-winged fly is speculated to result from the segmental identity shifting from the third to the second abdominal segment. *Ubx* has also been reported to determine the hindwing color pattern of adult butterflies (Matsuoka and Monteiro 2021; Tendolkar et al. 2021; Matsuoka and Monteiro 2022), and in this study, we found for the first time that *Ubx* is also involved in the color pattern of butterfly larvae, confirming that *Ubx* is co-opted to many traits in insects (Monteiro 2012). In addition, *Abd-B* repressed the expression of abd-A in the eighth and ninth abdominal segments of *Drosophila* embryos when they are present in the same cell (Karch et al. 1990). The RNAi experiment of *abd-A* and *Abd-B* in *P. memnon* resulted in the complete elimination and duplication of the V-shaped marking on the fifth and sixth abdominal segments (supplementary fig. S14). These results seem to suggest the possible segmental identity transformation after the knockdown of homeobox genes in *Papilio* caterpillars. As reported in Jin et al. (2019), only RNAi of prepatterning genes within the JH-sensitive period caused the phenotypic change. It is probably because the interaction between the JH signaling pathway and homeotic genes allows redefining the body plan, causing the color pattern switch in *Papilio* caterpillars.

## Materials and Methods

### Butterfly rearing

*P. memnon* females were collected on Amami-Ohshima Is., Kagoshima, Japan in August, 2020. We breed the mother butterflies to lay eggs on fresh citrus leaves in rectangular plastic containers with a size of 30×18×18 cm^3^, at 25 ℃, and under a 16:8 light: dark cycle. The eggs were collected in a round plastic cup (diameter, 100 mm; height, 45 mm) and the hatched caterpillars were fed on fresh citrus leaves until pupation and adult emergence in the rectangular plastic containers at 25 ℃ and with a 16:8 light: dark cycle.

### Juvenile hormone (JH) analog (fenoxycarb) treatment

JH analog application experiments were conducted following Futahashi and Fujiwara (2008) to ascertain the function of JH in switching larval patterns from 4th to 5th instar larvae in *P. memnon*. Fenoxycarb (Cayman Chemical Company, Michigan, USA), which serves as a carbamate insect growth regulator by mimicking juvenile hormone (JH), was dissolved in acetone (Wako Pure Chemical Industries, Japan). 5 μl of 1μg/μl fenoxycarb was ectopically applied to the larval body at the first 4 h, 12 h and 24 h during the penultimate (4th) instar. Larvae in the control group were treated with 5 μl acetone only. Larval developmental stage was confirmed by a camera-recording system with a 5-min capture interval (Cosmosoft Technologies, India). JH-analog-treated individuals were starved for 24 h and then reared in an independent incubator at 25 ℃ for further phenotypic observation.

### RNA sequencing (RNA-seq)

RNA-seq was performed to search for genes involved in larval green formation and larval pattern and to examine their expression levels. The anterior and posterior epidermal regions of third-, fourth-, and fifth-instar larvae, respectively, were sampled and RNA was extracted (fig. 1A). Sampling was done 12 hours after molting (fig. 1B). After quick anesthesia of caterpillars through a 5-min ice incubation, the caterpillars were soaked in the pre-cooled phosphate-buffered saline (PBS) and fixed on the wax dish by insect pins. Then gut, fat body, muscles and other accessory tissues were removed carefully without damaging the epidermis under a stereo microscope (Stemi 35, ZEISS). The total RNA was extracted using TRI reagent (Sigma) and treated by DNase I (TaKaRa, Japan). The total RNAs were sent to Macrogen Japan Corp. for paired-end, 101 bp transcriptomic sequencing by the illumina sequencer (NovaSeq6000). The RNA libraries were constructed using SMART-Seq^TM^ v4 Ultra^TM^ Low Input RNA Kit and TruSeq RNA Sample Prep Kit v2 by following the SMART-Seq^TM^ v4 Ultra^TM^ Low Input RN protocol. Given the limited size of epidermal samples, three individuals are mixed to ensure the sequencing depth.

Quality control checks of raw sequence read were performed by FastQC Version 0.11.9 (Andrews 2010). The raw sequence reads were mapped to the reference genome of *P. memnon* (BioProject accession ID: PRJDB5519) using HISAT2 v2.2.1 (Kim et al. 2019). The Sam file output from HISAT2 was sorted and converted to a BAM format using SAMtools v1.4 (Danecek et al. 2021). Potential full-length transcripts representing multiple splice variants for each gene locus were assembled and quantitated by StringTie v2.1.5 (Pertea et al. 2015). A reference annotation file was generated after assembling each RNA-seq library. Then all generated annotation files are merged to assemble the transfrags from the input annotation file with the reference transcripts, generating a global and non-redundant set of transcripts across multiple RNA-seq samples. The merged file of reference transcripts was used as the final annotation file for further analysis. Fragments Per Kilobase of transcript per Million mapped reads (FPKM) of each transcript was then calculated. Differentially expressed genes (DEGs) between two pairwise RNA-seq libraries (from different region, but at the same stage) were screened by edgeR v3.14 (Robinson et al. 2010; McCarthy et al. 2012; Chen et al. 2016) on R platform v4.1.0 (R Core Team). *p* value of each transcript was calculated by Fisher’s exact test using a hypothetical dispersion value, 0.1. Transcripts with a *p* value smaller than 0.05 and an absolute value of the transformed *Fold Change* greater than 1 were selected as DEGs, which were identified by Blastx (Camacho et al. 2009) against the Uniprot protein database (https://www.uniprot.org/downloads). The expression levels in 4th instar larvae of *P. xuthus* were quantified as described above by mapping to the reference genome of *P. xuthus* (BioProject accession ID: PRJNA291600, PRJDB2956) using RNA-seq data from a previous study (Jin et al., 2019).

### Identification of *JHBP* and *BBP* genetic cluster

Since the results of RNA-seq and previous studies (Futahashi et al. 2012; Yoda et al. 2020) suggested that *BBP* and *JHBP* are involved in larval green coloration of *Papilio* butterflies, *BBP* and *JHBP* clusters in *P. memnon* were identified. Four conserved neighboring genes in *P. polytes* [uncharacterized LOC106101173 (XM_013280315.1), hemicentin-1-like (XM_013280340.1), uncharacterized LOC106101173 (XM_013280315.1) and hemicentin-1-like (XM_013280340.1)] were used as indicator to search for the orthologous scaffolds of *BBP* cluster in *P. memnon*. With the four neighboring genes, a blastp program was performed against the genome of *P. memnon* (BioProject accession ID: PRJDB5519) and then the corresponding scaffold was retrieved. To identify the *BBP* cluster in *P. memnon*, the *BBP* orthologs of *P. polytes* were mapped to the retrieved scaffold (LD700013.1: dsx_scaffold_mimetic) using Geneious Prime v2021.2. Similarly, We used the *toll-like gene* in *P. polytes* (XM_013282841.1) as a landmark to indicate the genomic location of *JHBP* cluster and then mapped all the reported *JHBP*s in *P. polytes* to the corresponding orthologous scaffold (LD700011.1: chr_23homologous_long_hetro_left side). Subsequently, to confirm the reliability of the mapping results, phylogenetic analyses of multiple *BBP*s and *JHBP*s from five *Papilio* species (*P. memnon*, *P. polytes*, *P. xuthus*, *P. machaon* and *P. glaucus*) were performed by MEGA v10.1.7 (Kumar et al. 2018). The orthologous sequences of *BBP*s and *JHBP*s were aligned using multiple sequence alignment by ClustalW (Thompson et al. 1994). The phylogenetic tree was constructed by neighbor-joining method with an assessment of the confidence levels for various phylogenetic lineages using a 1000-replicated bootstrap analysis. Only when the results of the phylogenetic analyses were consistent with the mapping results, the orthologs of multiple *BBP*s and *JHBP*s in *P. memnon* were considered reliable.

### *in vivo* electroporation-mediated RNA interference

Efficient and target-specific short interfering RNA (siRNA) design was performed on the web server siDirect 2.0 (http://sidirect2.rnai.jp/). G/C content of target sequence was subject to 35%∼ 55%. Contiguous G’s or C’s 4 nt or more are avoided for easier chemical synthesis of siRNA. A maximum Tm value in seed sequence was set to 21.5 ℃ to minimize the seed-dependent off-target effects (Ui-Tei et al. 2008). siRNAs were purchased through Custom oligonucleotide Synthesis service provided by Food Assessment & Management Center (FASMAC Co. Ltd., Atsugi, Japan). Dry powder siRNA was dissolved in the annealing buffer [100 mM CH3COOK, 2 mM Mg (CH_3_COO)_2_, 30 mM HEPES-KOH, pH 7.4] to a concentration at 500 μM and then stored at -80 ℃. Sequence information of siRNA is listed in Supplementary table S3.

Functional analysis of candidate genes was performed using *in vivo* electroporation-mediated RNA interference (Ando and Fujiwara 2013). Injection capillary, processed on versatile puller (Narishige Group, Tokyo, Japan) by automated double pulling, was prepared from ultrasonic cleaned glass capillary with a filament (Narishige Group, Tokyo, Japan). siRNA was loaded into the injection capillary and injected through an electronic microinjector FemtoJet® (Eppendorf AG., Tokyo, Japan). *Papilio* caterpillar was gently fixed on an operational platform by tape. The position of capillary was controlled by a three-dimensional manual micromanipulator (Narishige Group, Tokyo, Japan). 1 μl of 250 μM siRNA was then injected into the hemolymph of caterpillars through the intersegmental membrane between 7th abdominal segment and 8th abdominal segment. Two PBS droplets were placed as buffering conductor between electrodes and the larval cuticle to reduce damage to epidermis. To incorporate the siRNA into the target epidermal cells, electroporation with a poring pulse and a subsequent transfer pulse was conducted by a super electroporator (Nepa Gene Co., Ltd., Tokyo, Japan). Detailed conditions of electroporation are listed in Supplementary table S5. Universal negative control siRNA (Nippon Gene) was used as a negative control (supplementary fig. S18).

## Acknowledgements

We thank Dr. H. Jin for helpful comments and experimental supports.

## Data Availability

The datasets presented in this study can be found in online repositories. All generated sequence data are available from National Center for Biotechnology Information (BioProject: PRJNA964473, https://dataview.ncbi.nlm.nih.gov/object/PRJNA964473?reviewer=cfurjocqicb0kn66n0in5st3lg)

## Conflict of Interest

The authors declare that the research was conducted in the absence of any commercial or financial relationships that could be construed as a potential conflict of interest.

## Author Contributions

LL and HF conceived and designed the study. LL, SK and KW conducted experiments. LL, SK, TK and HF wrote the paper. HF supervised this project. All authors reviewed the manuscript.

## Funding

This work was supported by Ministry of Education, Culture, Sports, Science and Technology/Japan Society for the Promotion of Science KAKENHI (22128005, 15H05778, 18H04880, 20H04918, 20H00474 to HF; 19J00715 to SK), JST SPRING (Grant Number: JPMJSP2108 to LL) and China Scholarship Council (CSC, 202008360145 to KW).

## Notes

### Competing Interest Statement

The authors have declared no competing interest.

