## Supplemental materials for "Molecular mechanisms underlying the formation of larval green color and camouflage patterns in swallowtail butterfly, *Papilio memnon*"

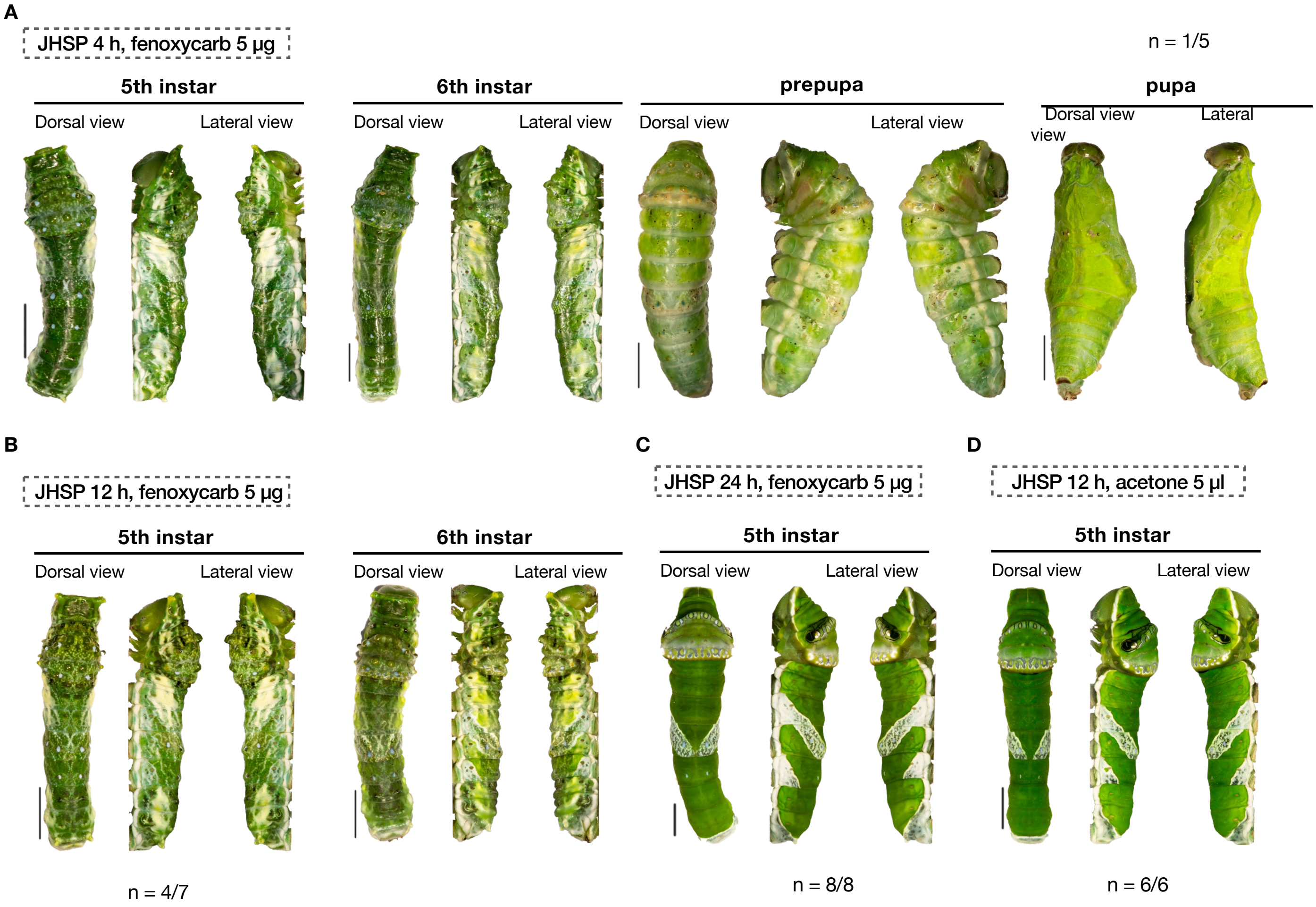


**Figure S1. Juvenile Hormone-analog (JHA) treatment in *P. memnon*.** (A) Individuals treated with 5 µg fenoxycarb at 4 h after the third molt. 4/5 of the individuals did not survive till the 5th instar. (B) Individuals treated with 5 µg fenoxycarb at 12 h after the third molt. 2/9 of the individuals did not survive till the 5th instar. (C) Individuals treated with 5 µg fenoxycarb at 24 h after the third molt. (D) Individuals treated with 5 µg acetone at 12 h after the third molt. JHSP: Juvenile hormone sensitive period. The color pattern switch (from mimetic to camouflage) was inhibited by the JHA treatment during JHSP, generating an additional instar.

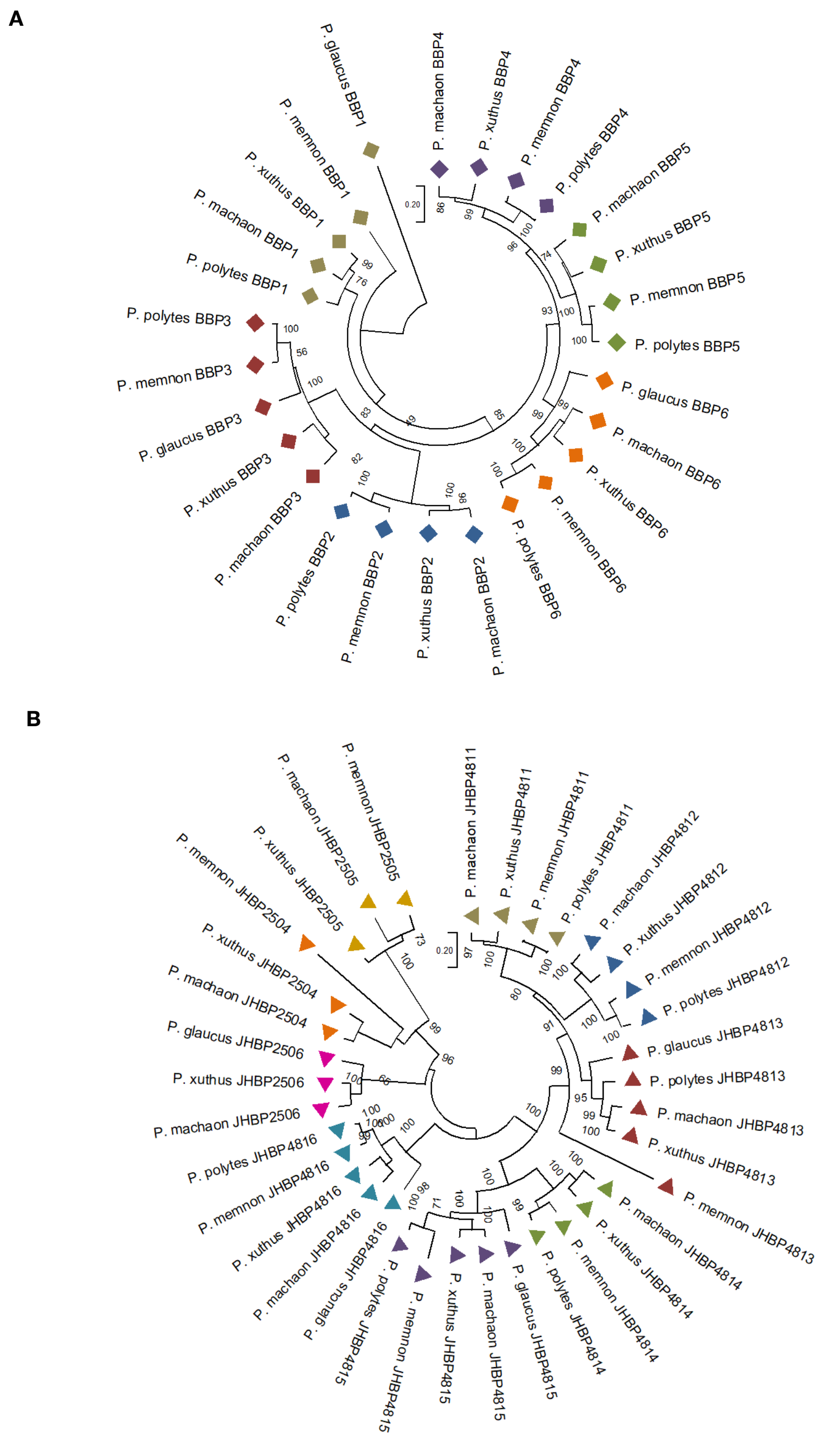


**Figure S2. Phylogenetic tree of *bilin-binding proteins* (*BBP*s) and *JH-binding proteins* (*JHBP*s).** Unrooted neighbor-joining tree of *BBP*s homologs. The Khaki, blue, reddish-brown, purple, green and orange squares indicate the homologs of *BBP1*, *BBP2*, *BBP3*, *BBP4*, *BBP5* and *BBP6*, respectively. Bootstraps values are shown at the tree nodes. *P. xuthus*: *Papilio xuthus*. *P. polytes*: *Papilio polytes*. *P. memnon*: *Papilio memnon*. *P. machaon*: *Papilio machaon*. *P. glaucus*: *Papilio glaucus*. (B) Unrooted neighbor-joining tree of *JHBP*s homologs. The Khaki, blue, reddish-brown, green, purple, cyan, orange, yellow and pink triangles indicate the homologs of *JHBP4811*, *JHBP4812*, *JHBP4813*, *JHBP4814*, *JHBP4815, JHBP4816*, *JHBP2504*, *JHBP2505* and *JHBP2506*, respectively. Bootstraps values are shown at the tree nodes. *P. xuthus*: *Papilio xuthus*. *P. polytes*: *Papilio polytes*. *P. memnon*: *Papilio memnon*. *P. machaon*: *Papilio machaon*. *P. glaucus*: *Papilio glaucus*.

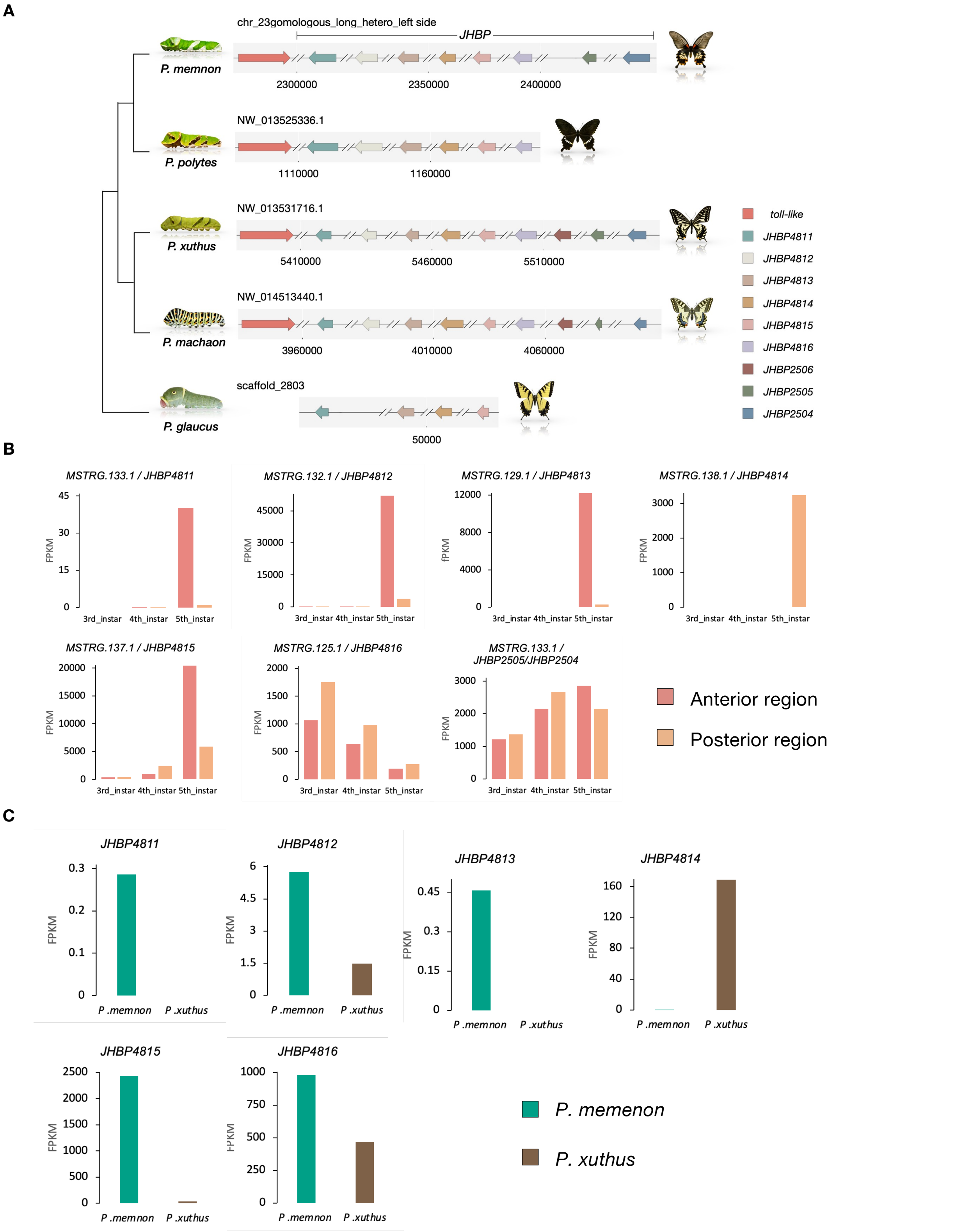


**Figure S3. Structure and expression profiles of *JH-binding proteins* (*JHBPs*).** (A) Cluster of *JHBP*s in 5 *Papilio* species. The length of arrows represents the relative gene length. Transcriptional orientation is shown by the direction of arrows. (B) Expression profiles of *JHBP*s from 3rd- to 5th-instar in epidemics. (C) Epidermal expression of *JHBP*s at 4th-instar in *P. memnon*. (RNAseq data, n =1 but one sample is mixed with three individuals). The expression level of *JHBP*s is shown by the fragments per kilobase of exon per million mapped fragments (FPKM).

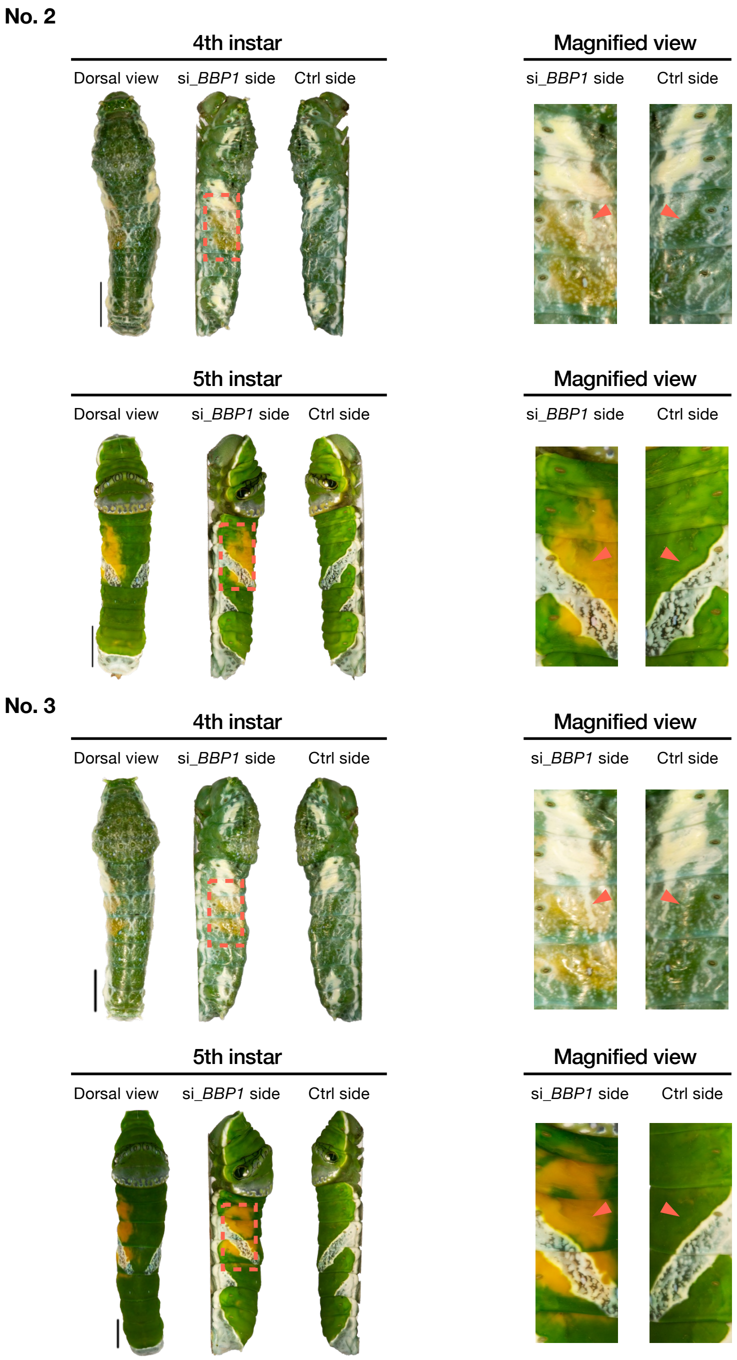


**Figure S4. in vivo electroporation-mediated RNAi of *BBP1*.** Additional individuals. 4th-instar (left) and 5th-instar (right) larvae after RNAi of *BBP1* siRNA is injected through the intersegmental membrane between the 7th and 8th abdominal segment and introduced into specific epidermal region (indicated by red dashed frame) via an electroporation-mediated method during the 3rd-instar. scale bar = 5 mm.

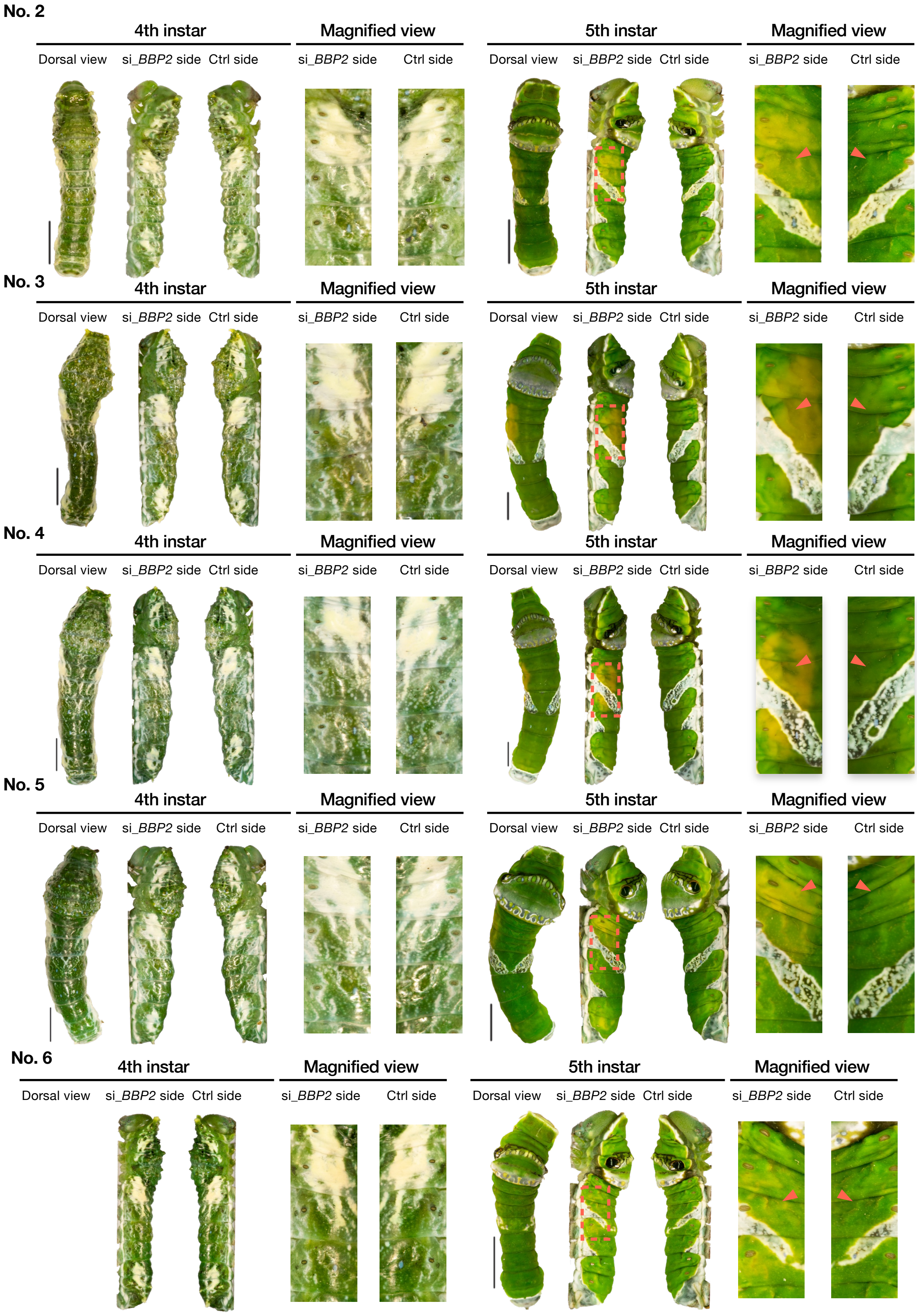


**Figure S5. in vivo electroporation-mediated RNAi of BBP2.** Additional individuals. 4th-instar (left) and 5th-instar (right) larvae after RNAi of BBP2 siRNA is injected through the intersegmental membrane between the 7th and 8th abdominal segment and introduced into specific epidermal region (indicated by red dashed frame) via an electroporation-mediated method during the 3rd-instar. scale bar = 5 mm.

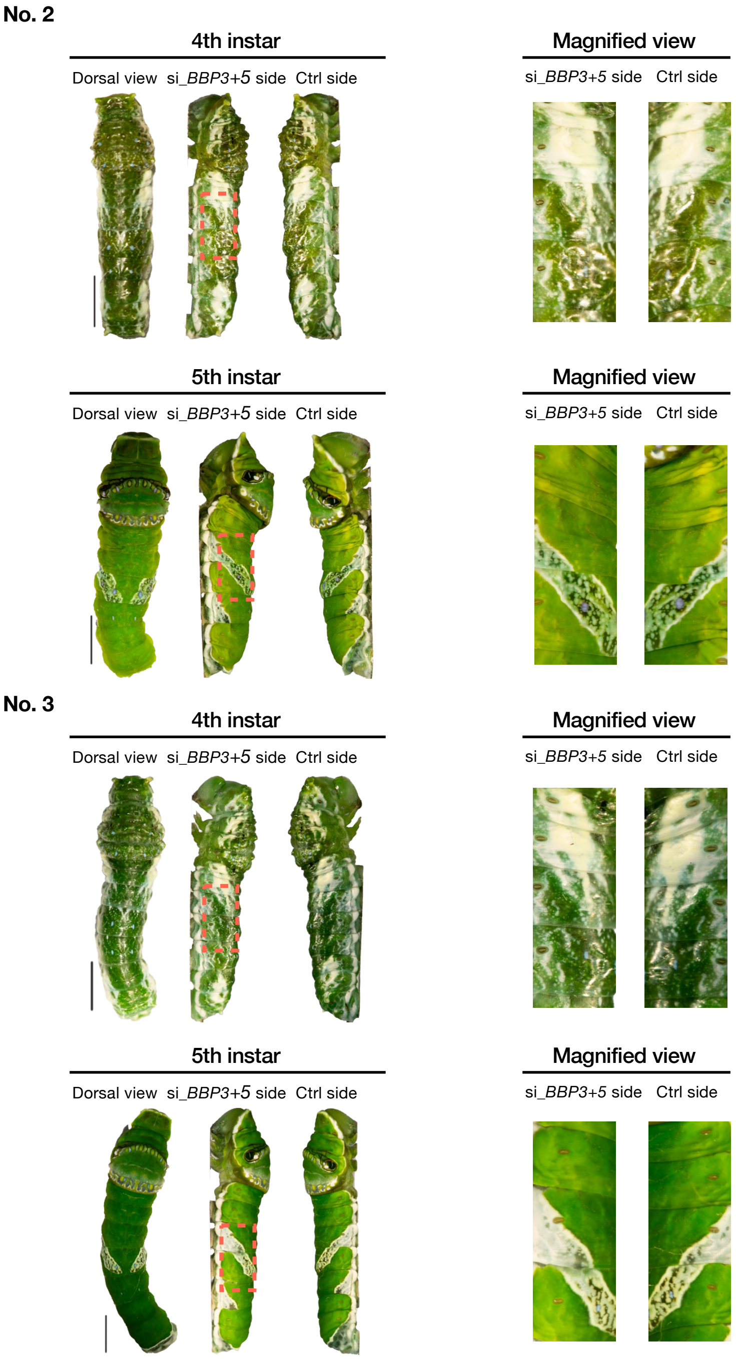


**Figure S6. in vivo electroporation-mediated RNAi of *BBP 3* + *5*.** Additional individuals. 4th-instar (left) and 5th-instar (right) larvae after RNAi of *BBP3* + *5* siRNA is injected through the intersegmental membrane between the 7th and 8th abdominal segment and introduced into specific epidermal region (indicated by red dashed frame) via an electroporation-mediated method during the 3rd-instar. scale bar = 5 mm.

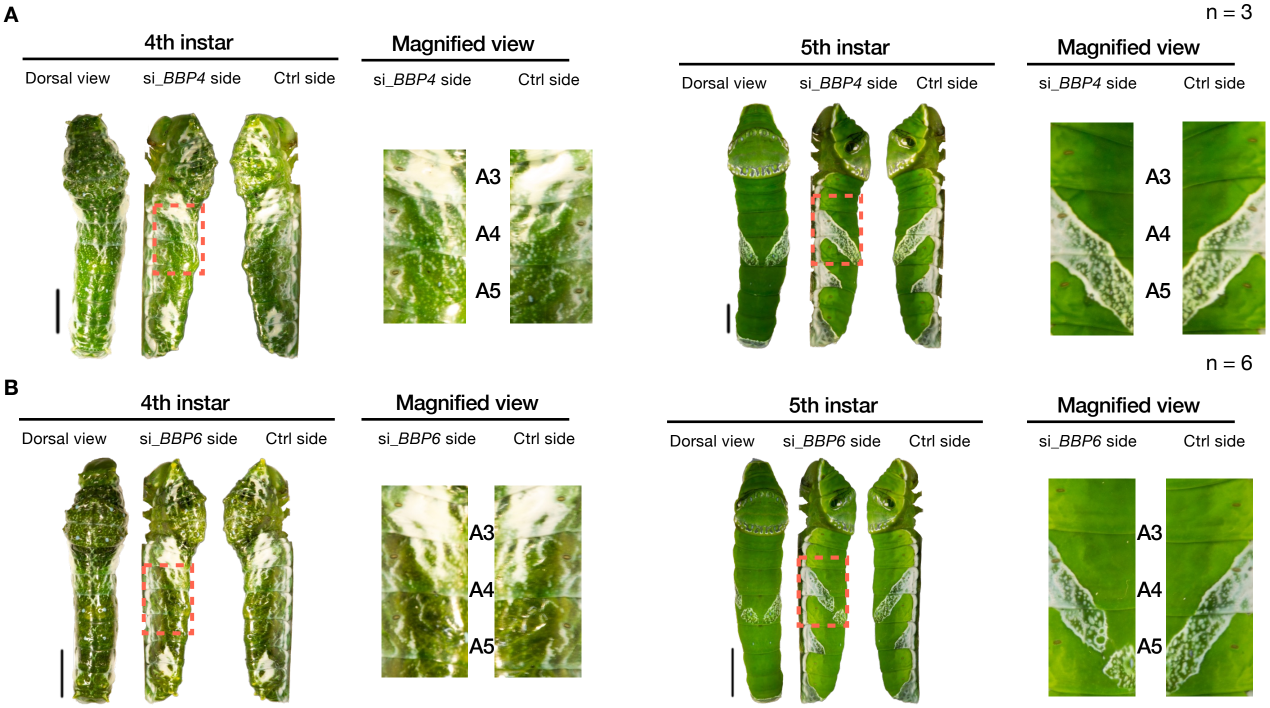


**Figure S7. in vivo electroporation-mediated RNAi of *BB4* and *BBP6*.** (A) 4th-instar (left) and 5th-instar (right) larvae after RNAi of *BBP4* (B) 4th-instar (left) and 5th-instar (right) larvae after RNAi of *BBP6.* siRNA is injected through the intersegmental membrane between the 7th and 8th abdominal segment and introduced into specific epidermal region (indicated by red dashed frame) via an electroporation-mediated method during the 3rd-insar. scale bar = 5 mm. The knockdown of *BBP6* resulted in the elimination of the V-shaped pattern across the 4th and 5th abdomianl segments.

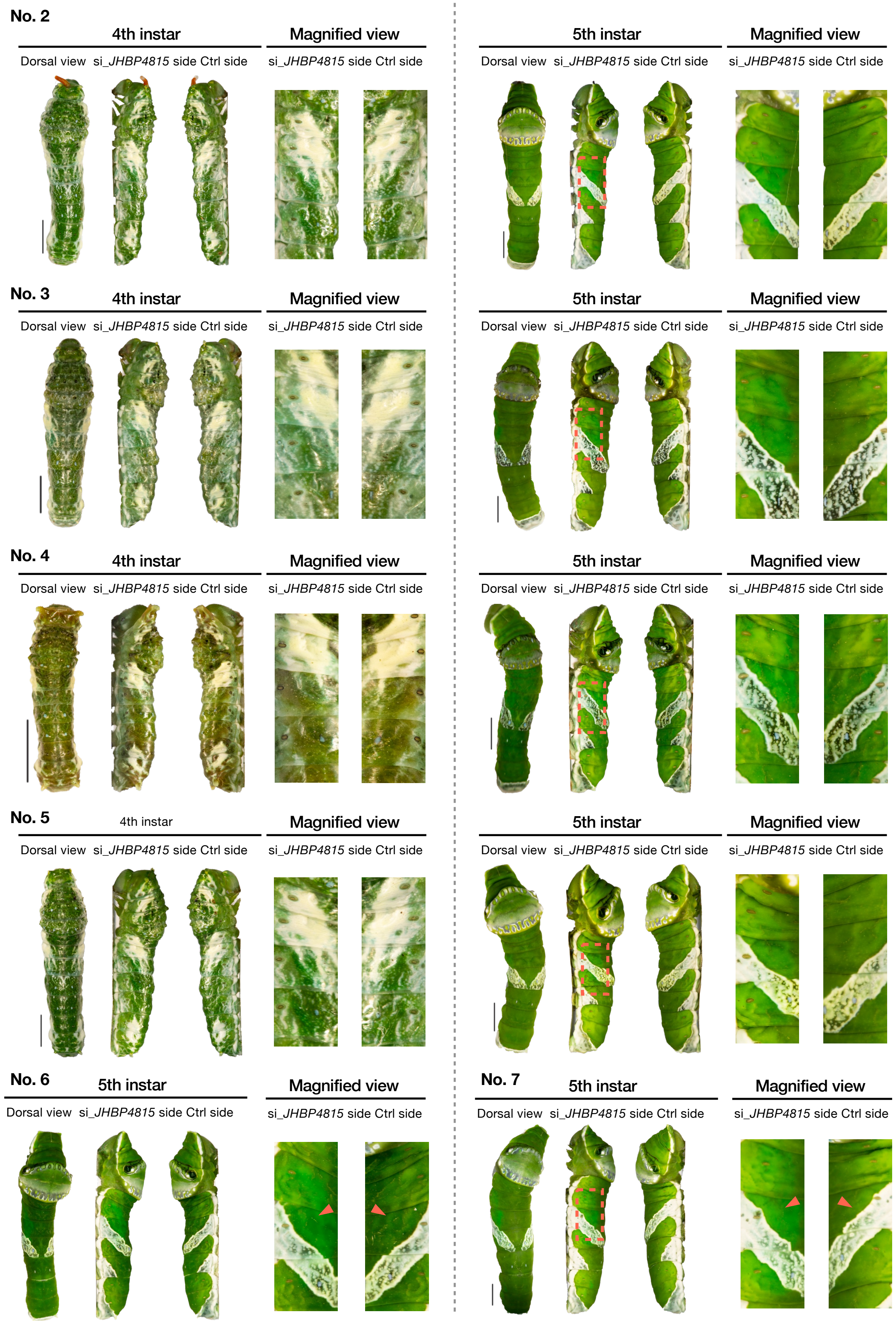


**Figure S8. in vivo electroporation-mediated RNAi of *JHBP4815*.** Additional individuals. 4th-instar (left) and 5th-instar (right) larvae after RNAi of *JHBP4815*. siRNA is injected through the intersegmental membrane between the 7th and 8th abdominal segment and introduced into specific epidermal region (indicated by red dashed frame) via an electroporation-mediated method during the 3rd-insar. scale bar = 5 mm.

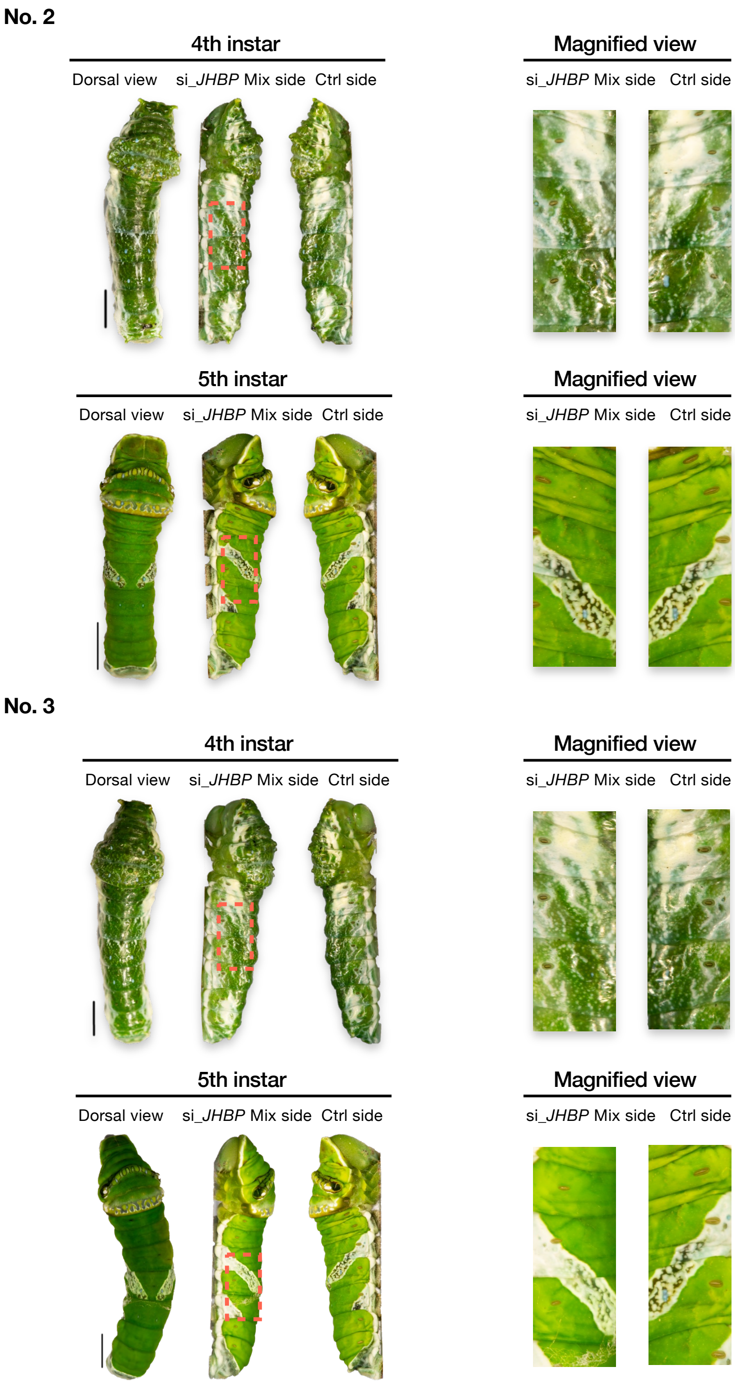


**Figure S9. in vivo electroporation-mediated RNAi of *JHBP*s.** Additional individuals. 4th-instar (left) and 5th-instar (right) larvae after multiple-RNAi of *JHBP4812, JHBP4814, JHBP4816 and JHBP2505.* siRNA is injected through the intersegmental membrane between the 7th and 8th abdominal segment and introduced into specific epidermal region (indicated by red dashed frame) via an electroporation-mediated method during the 3rd-insar. scale bar = 5 mm. Among all the RNAi experiments targeting *JHBP*s, only knockdown of *JHBP4815* caused a slight reduction of yellow color during the larval stage.

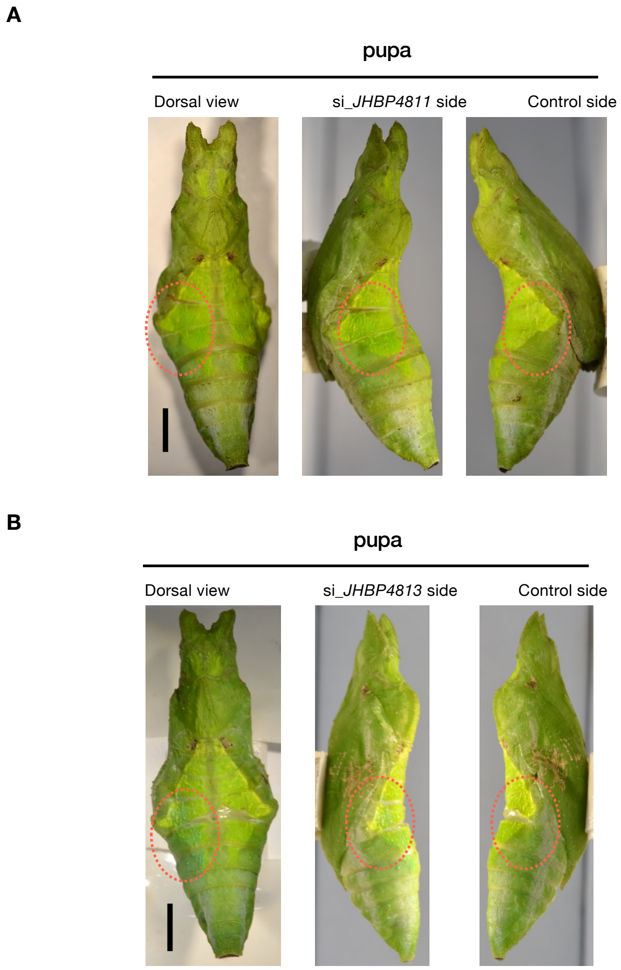


**Figure S10. Electroporation-mediated RNAi of *JHBP4811* and *JHBP4813*.** (A) pupa after the RNAi of *JHBP4811, n* = 4. (B) pupa after the RNAi of *JHBP4813, n* = 2. siRNA is injected through the intersegmental membrane between the 7th and 8th abdominal segment, and introduced into specific epidermal region (indicated by red dashed frame) via an electroporation-mediated method. scale bar = 5 mm.

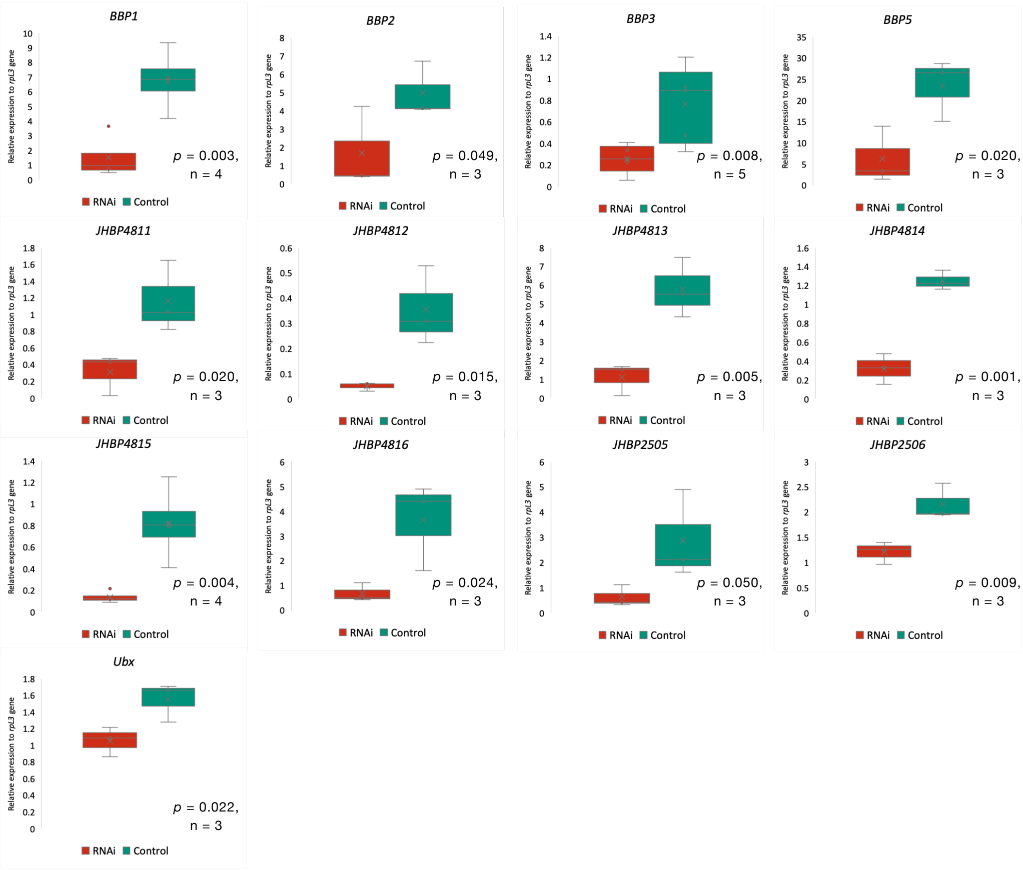


**Figure S11. Knockdown efficiency of RNAi experiments from qPCR.** *ribosomal protein L3* (*rpL3*) was used as a reference gene. Relative expressions of the RNAi and control sides were plotted in respective red and green colors. Student's t-test was performed to obtain the statistical significance. Primers used in this study are shown in Table S4.

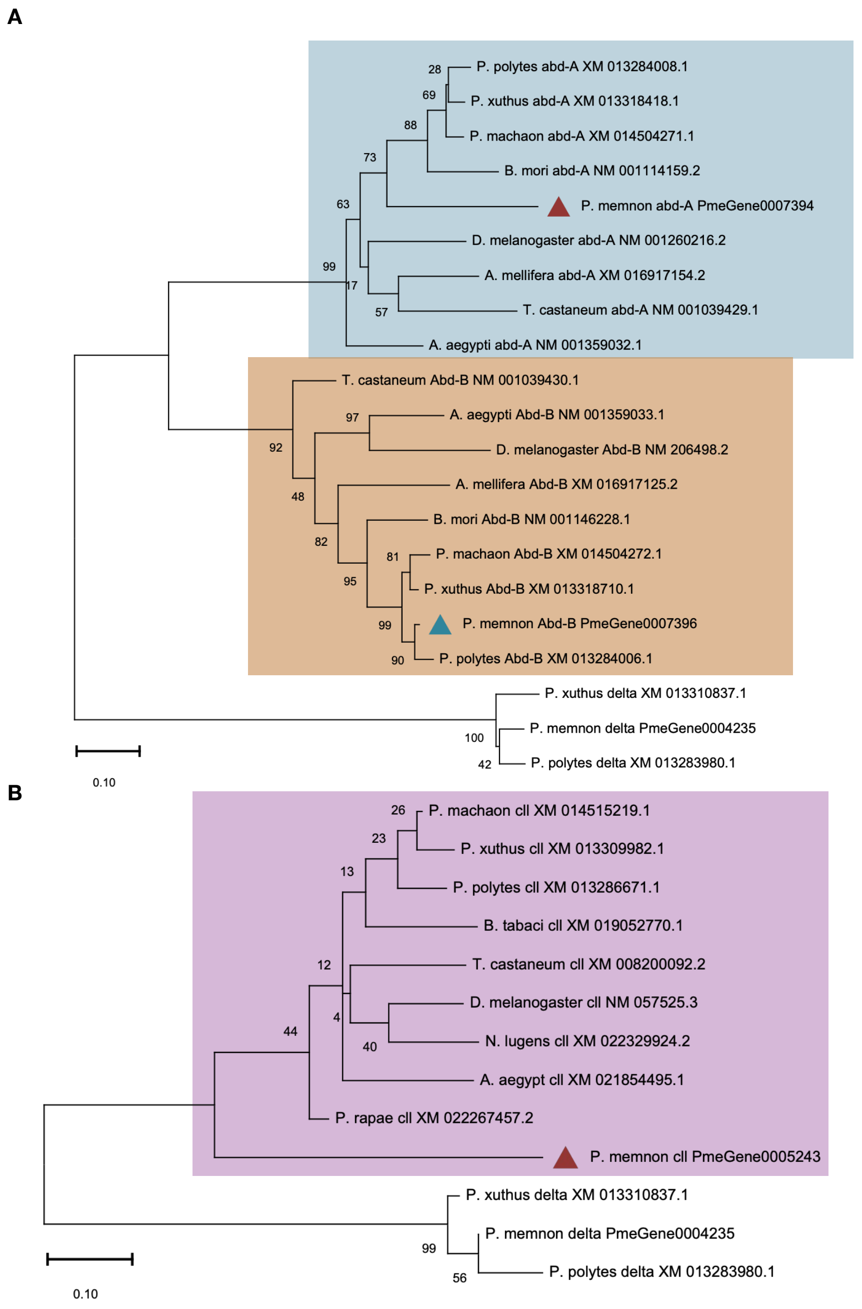


**Figure S12.** Phylogenetic tree of *clawless* (*cll*), *abdominal-A* (*abd-A*) and *abdominal-B* (*Abd-B*). Bootstrap values are shown at the tree nodes. *P. xuthus*: *Papilio xuthus. P. polytes*: *Papilio polytes. P. memnon*: *Papilio memnon. P. machaon*: *Papilio machaon*. *P. glaucus*: *Papilio glaucus*.

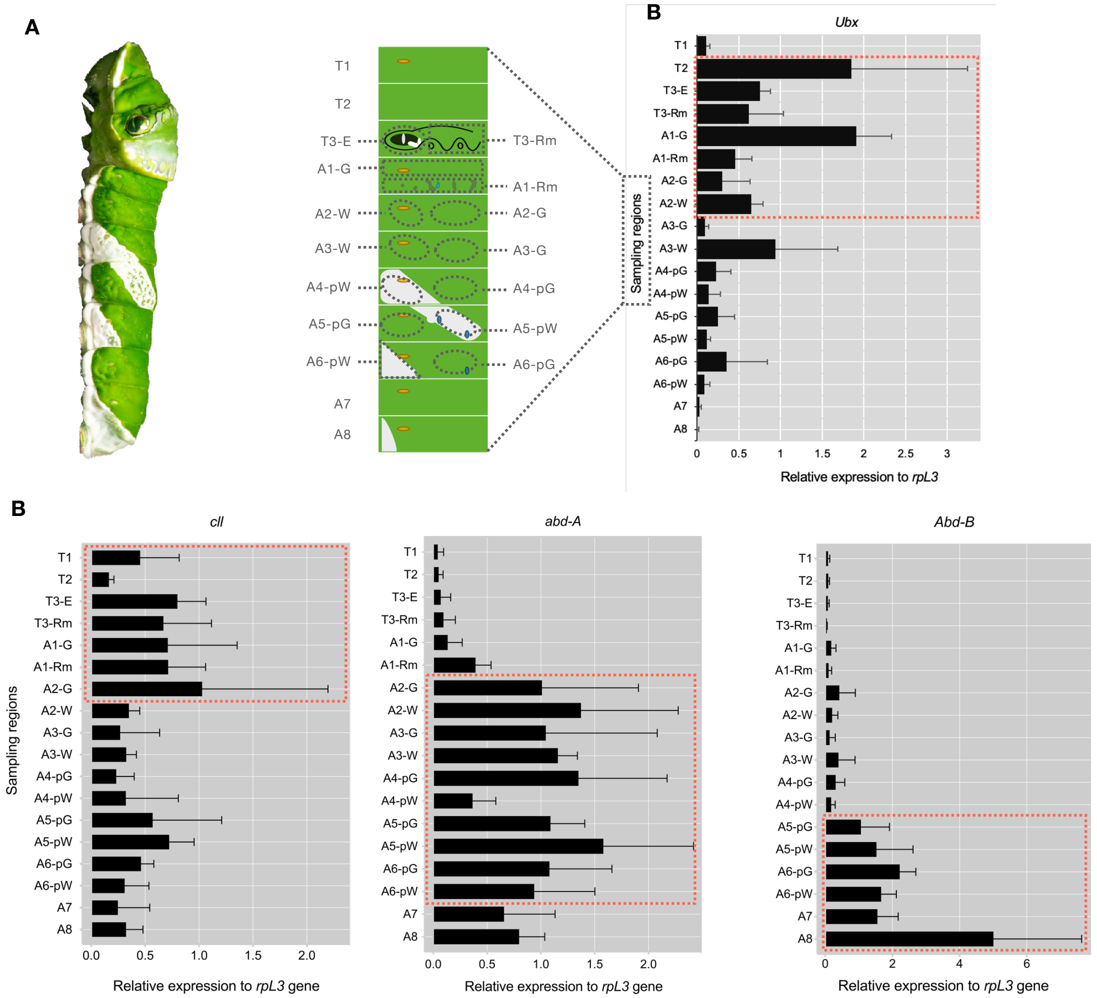


**Figure S13. Spatial expression pattern of prepatterning genes during Jh-sensitive period (JHSP) in *P. memnon*.** (A) Sampling region of the epidermal samples. (B) Relative expression of *Ubx*, *cll*, *abd-A* and *Abd-B* using the corresponding epidermal samples as shown in (A). Expression levels of target genes were tested by quantitative RT-PCR as described in Jin et al., 2019. The *ribosomal protein L3* gene (*rpL3*) was used as a reference gene for endogenous control. All data are presented as mean ± SD. The red dashed frame indicates where the expression is concentrated. n = 3.

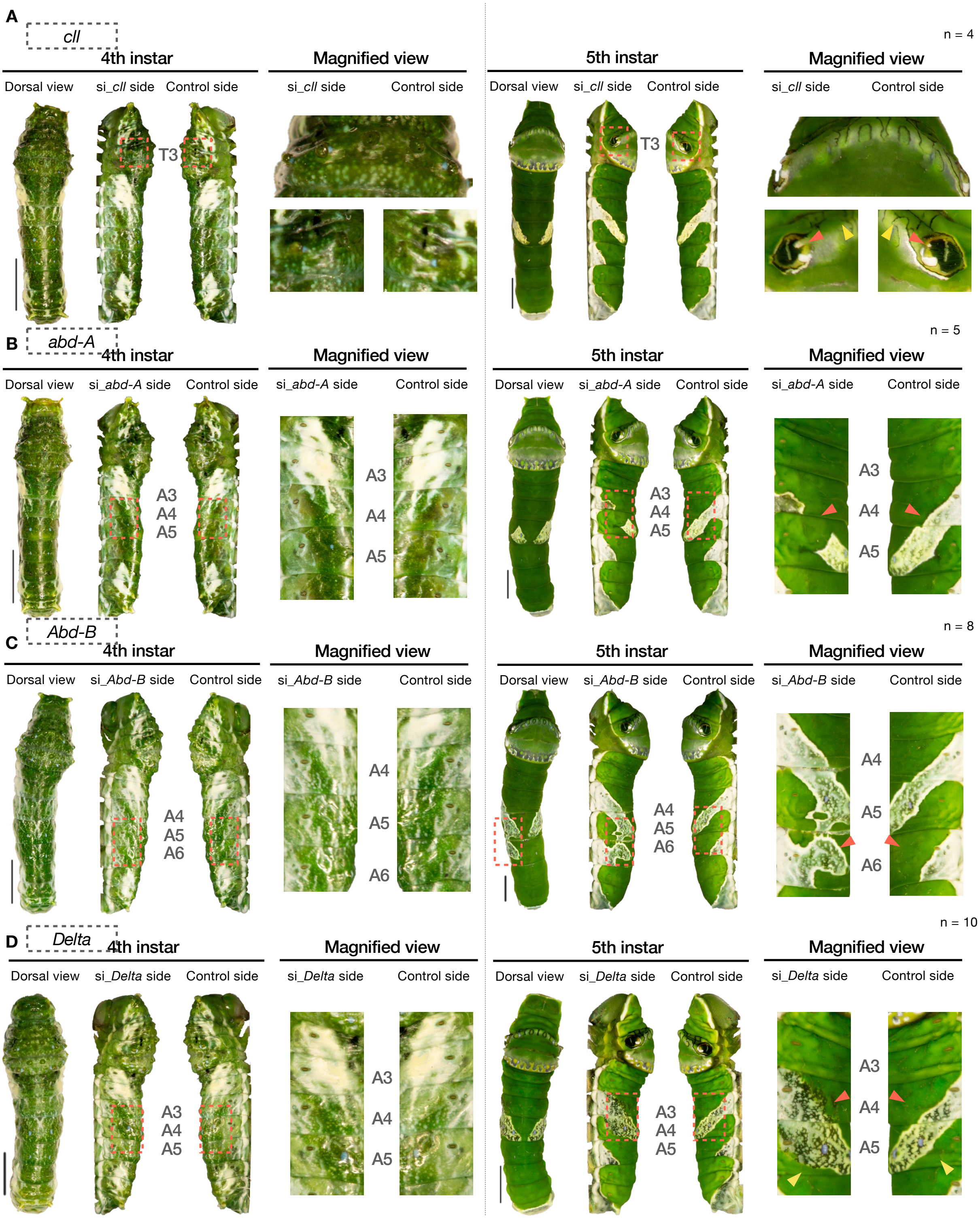


**Figure S14. Electroporation-mediated RNAi of *cll, abd-A, Abd-B* and *delta*.** (A) 4th-insatr (left) and 5th-instar larvae after RNAi of *cll* (B) 4th-insatr (left) and 5th-instar larvae after RNAi of *abd-A* (C) 4th-insatr (left) and 5th-instar larvae after RNAi of *Abd-B*. (D) 4th-insatr (left) and 5th-instar larvae after RNAi of *Delta*. siRNA is injected through the intersegmental membrane between the 7th and 8th abdominal segment, and introduced into specific epidermal region (indicated by red dashed frame) via an electroporation-mediated method. scale bar = 5 mm.

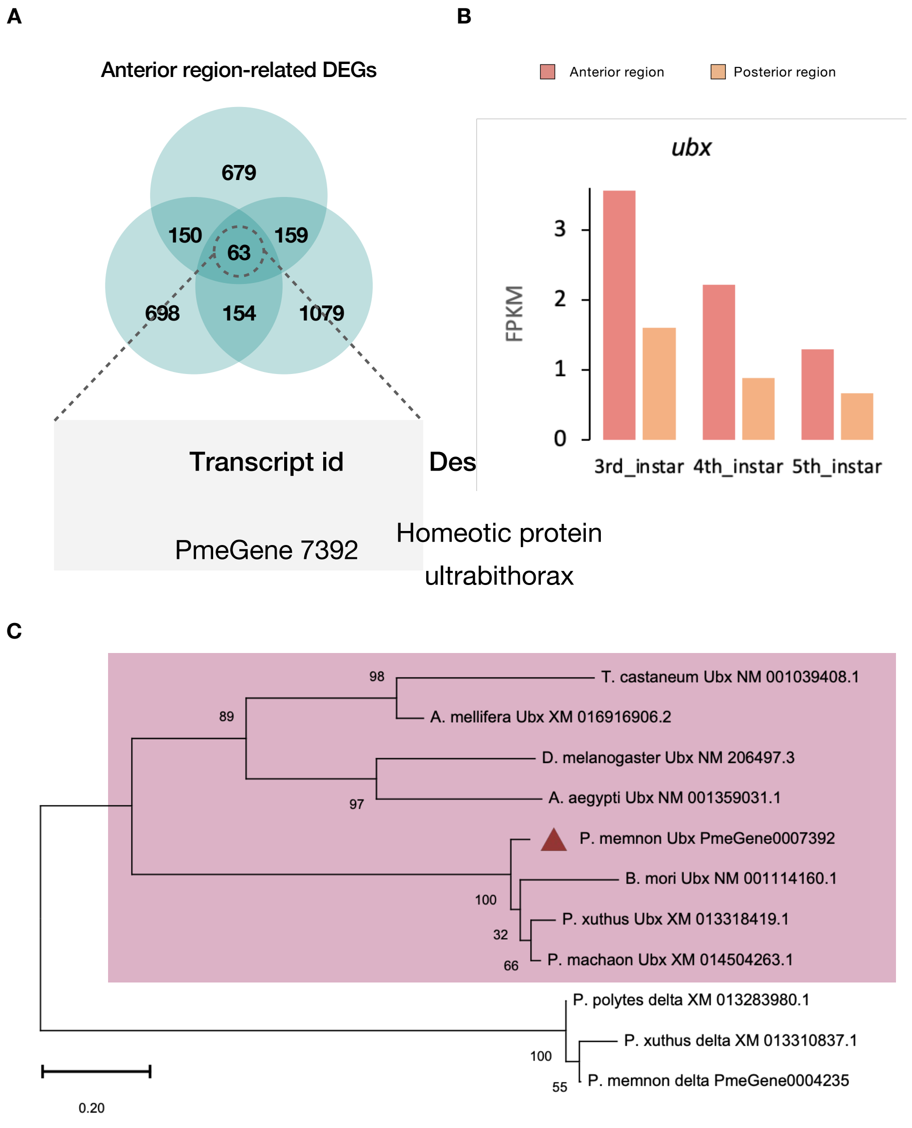


**Figure S15.** (A) Anterior region-associated Differentially Expressed Genes (DEGs). (B) Expression profiles of *Ultrabithorax* (*Ubx*) from 3rd- to 5th-instar in epidemics. (C) Phylogenetic tree of *Ubx.* Unrooted neighbor-joining tree of *BBP*s homologs. Bootstrap values are shown at the tree nodes. *P. xuthus*: *Papilio xuthus. P. polytes*: *Papilio polytes. P. memnon*: *Papilio memnon. P. machaon*: *Papilio machaon. P. glaucus*: *Papilio glaucus.*

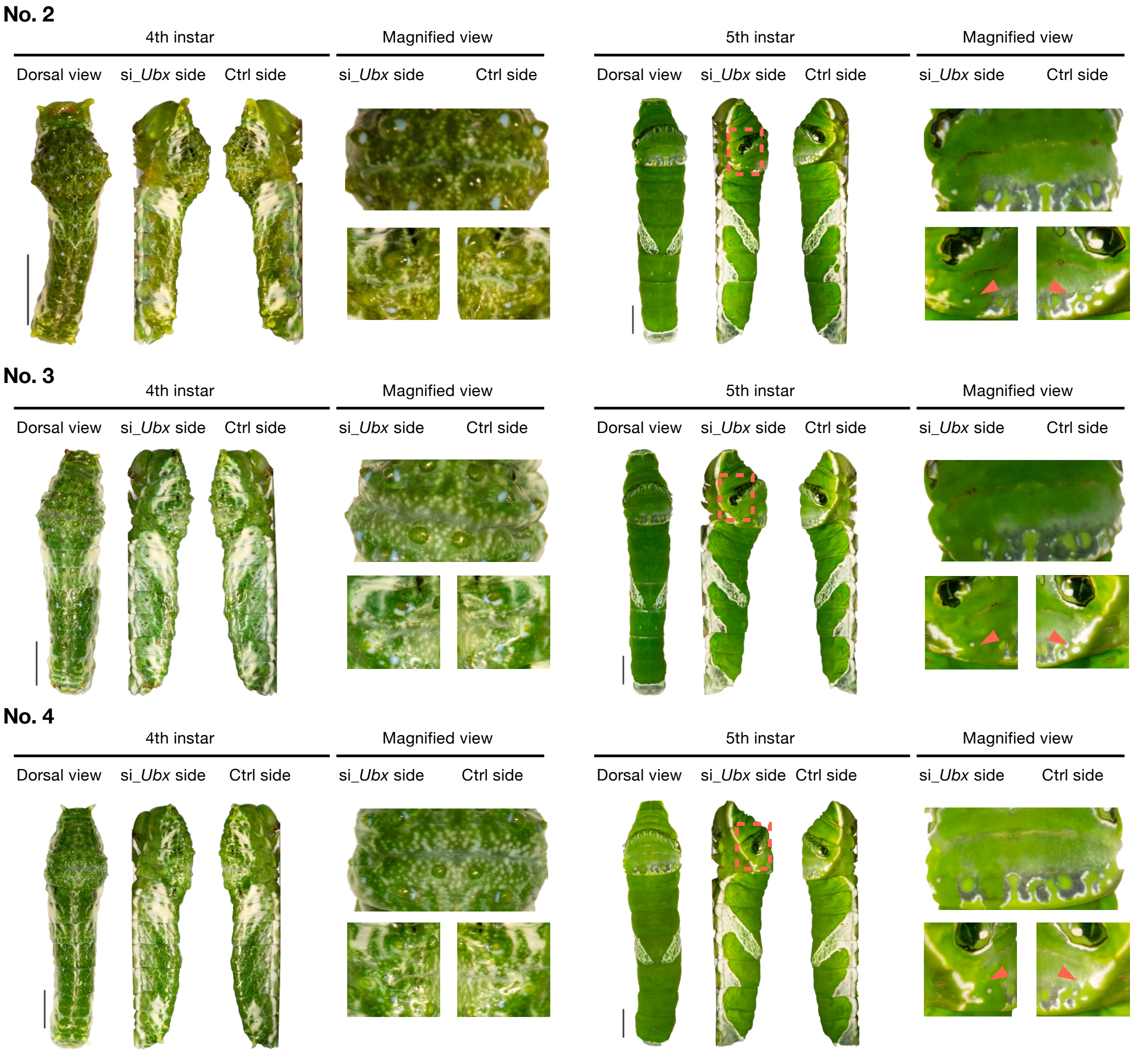


**Figure S16. in vivo electroporation-mediated RNAi of *Ubx* on the first abdominal se6ment (A1).** Additional individuals. *Ubx* knockdown on the first abdominal segment (A1, indicated by the red box). siRNA is injected through the intersegmental membrane between the 7th and 8th abdominal segment, and introduced into specific epidermal regions (indicated by red dashed frame) via an electroporation-mediated method during the 3rd-instar. scale bar = 5 mm.

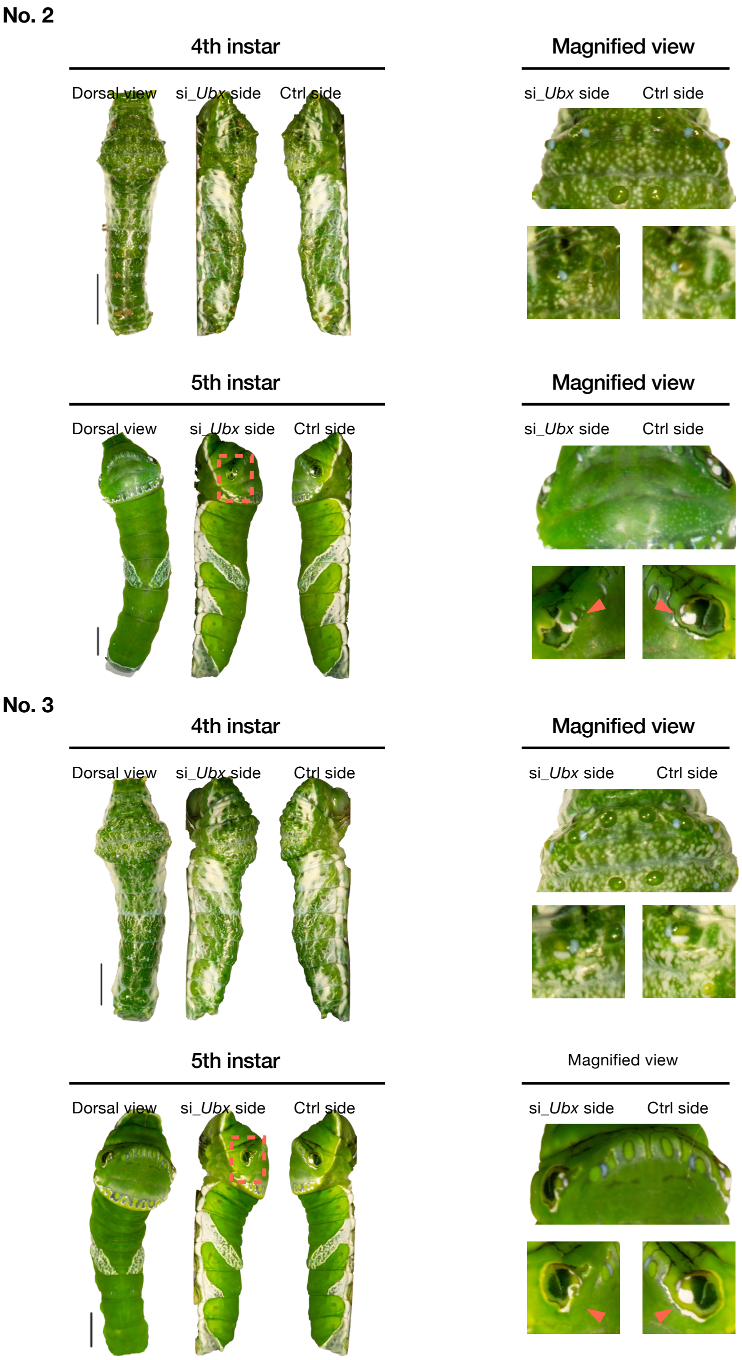


**Figure S17. in vivo electroporation-mediated RNAi of *Ubx* on the third thoracic segment (T3).** Additional individuals. *Ubx* knockdown on the first abdominal segment (A1, indicated by the red box). siRNA is injected through the intersegmental membrane between the 7th and 8th abdominal segment, and introduced into specific epidermal regions (indicated by red dashed frame) via an electroporation-mediated method during the 3rd-instar. scale bar = 5 mm.

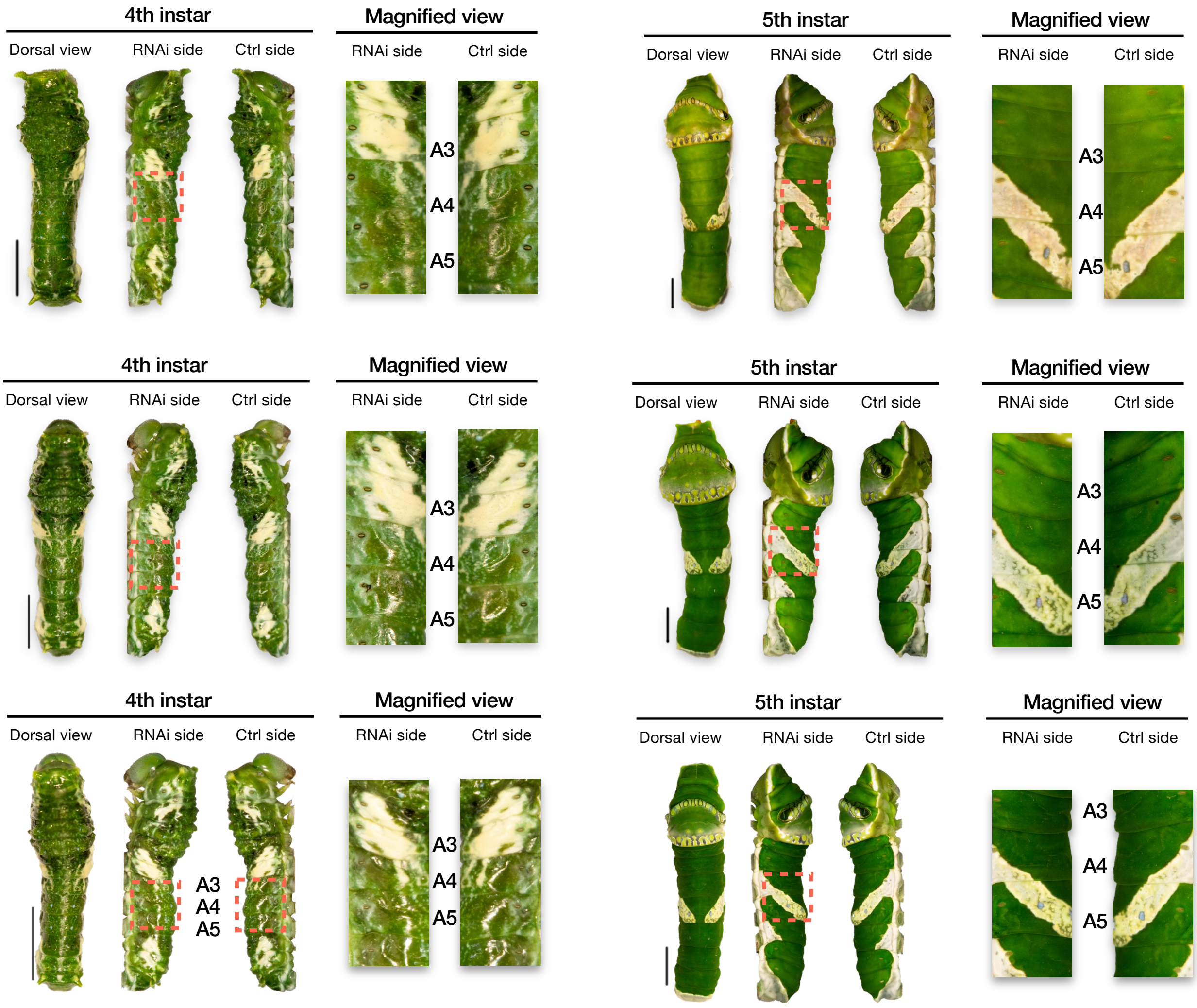


**Figure S18. Negative Control**. Left panel, the phenotype of the 4th instar larva after the electroporation. Right panel, the phenotype of the 5th instar larva after the electroporation. 1 μl 250 mM universal siRNA was injected at the late 3rd-instar stage. The red frames represent the region where the electroporation was performed. A3, A4 and A5: the third, fourth and fifth abdominal segments, respectively. scale bar=5 mm, n=3.

**Table S1. Statistics of RNA-seq raw data.**

| Sample name | read counts | read bases base pair (bp) | GC content |
|---|---|---|---|
| 3rd_A | 53,723,644 | 5,426,088,044 | 33.57% |
| 3rd_P | 50,693,406 | 5,120,034,006 | 39.56% |
| 4th_A | 54,697,212 | 5,524,418,412 | 35.8% |
| 4th_P | 43,657,636 | 4,409,421,236 | 37.7% |
| 5th_A | 41,849,038 | 41,849,038 | 42.07% |
| 5th_P | 45,115,144 | 4,556,629,544 | 41.32% |

**Table S2. Green color-associated DEGs.** Gens were manually annotated and categorized according to the local blast results.

| Transcripts ID | Assession No. | Description | PValue | FPKM (3rd_P) | PValue | FPKM (4th_P) | PValue | FPKM (5th_A) | Category |
|---|---|---|---|---|---|---|---|---|---|
| MSTRG.10015.1 | XM_013293875 | PREDICTED: Papilio polytes filaggrin-2-like (LOC106111732), mRNA | 0.000200699 | 1.509861 | 0.003182373 | 0.390062 | 6.71E-39 | 15.667977 | Cellular events-associated |
| MSTRG.11807.2 | NM_001311465 | Papilio polytes 40S ribosomal protein SA (LOC106107199), mRNA | 5.27E-38 | 73.057648 | 1.94E-44 | 50.648502 | 4.38E-11 | 83.081467 | Cellular events-associated |
| MSTRG.1795.5 | XM_013292307 | PREDICTED: Papilio polytes protein anoxia up-regulated-like (LOC106110465), transcript variant X4, mRNA | 8.45E-08 | 0.617438 | 1.02E-05 | 0.694145 | 2.44E-33 | 3.662393 | Cellular events-associated |
| MSTRG.3046.5 | XM_013282342 | PREDICTED: Papilio polytes synaptic vesicle glycoprotein 2B-like (LOC106102767), mRNA | 3.31E-33 | 5.44021 | 4.03E-06 | 0.524453 | 4.31E-17 | 2.219629 | Cellular events-associated |
| MSTRG.4073.3 | NM_001311702 | Papilio polytes 60S acidic ribosomal protein P2 (LOC106108361), mRNA | 4.69E-73 | 415.950684 | 1.24E-06 | 462.334198 | 8.50E-79 | 145.469482 | Cellular events-associated |
| MSTRG.5144.1 | XM_013282519 | PREDICTED: Papilio polytes proton-coupled amino acid transporter 4-like (LOC106102899), mRNA | 7.69E-12 | 7.391792 | 5.98E-74 | 3.265747 | 2.35E-61 | 2.244452 | Cellular events-associated |
| MSTRG.5425.2 | XM_013278201 | PREDICTED: Papilio polytes tRNA-dihydrouridine(47) synthase [NAD(P)(+)]-like (LOC106099612), mRNA | 7.11E-16 | 0.78487 | 1.04E-14 | 0.467885 | 5.03E-19 | 0.898446 | Cellular events-associated |
| MSTRG.6002.1 | XM_013281597 | PREDICTED: Papilio polytes lysM and putative peptidoglycan-binding domain-containing protein 3 (LOC106102184), transcript variant X2, mRNA | 0.003429501 | 0.832925 | 2.30E-08 | 0.783711 | 1.65E-18 | 1.839301 | Cellular events-associated |
| MSTRG.6071.1 | XM_013286268 | PREDICTED: Papilio polytes cytochrome P450 6B2-like (LOC106105813), transcript variant X3, mRNA | 9.90E-15 | 0.462777 | 7.04E-09 | 0.249237 | 2.39E-13 | 0.883102 | Cellular events-associated |
| MSTRG.6408.3 | XM_013285503 | PREDICTED: Papilio polytes PI-PLC X domain-containing protein 3 (LOC106105226), transcript variant X2, mRNA | 0.000115682 | 2.369963 | 1.84E-05 | 1.785494 | 1.70E-05 | 1.290411 | Cellular events-associated |
| MSTRG.6534.1 | XM_013287773 | PREDICTED: Papilio polytes acyl-CoA Delta(11) desaturase-like (LOC106107071), mRNA | 6.51E-05 | 0.663441 | 9.11E-25 | 2.016498 | 1.01E-10 | 4.862396 | Cellular events-associated |
| MSTRG.6553.1 | XM_013284933 | PREDICTED: Papilio polytes organic cation transporter protein-like (LOC106104795), mRNA | 8.59E-11 | 15.903947 | 4.28E-05 | 0.642804 | 1.06E-09 | 0.763624 | Cellular events-associated |
| MSTRG.6622.2 | XM_013284943 | PREDICTED: Papilio polytes cytochrome P450 6B1-like (LOC106104802), mRNA | 2.07E-05 | 1.017942 | 0.019169253 | 0.34264 | 0.025042572 | 0.235988 | Cellular events-associated |
| MSTRG.7533.1 | XM_013278984 | PREDICTED: Papilio polytes vesicular glutamate transporter 1 (LOC106100208), transcript variant X2, mRNA | 0.01534811 | 0.387437 | 0.000662286 | 2.103111 | 4.83E-06 | 6.272163 | Cellular events-associated |
| MSTRG.7659.7 | XM_013277821 | PREDICTED: Papilio polytes probable sulfite oxidase, mitochondrial (LOC106099309), transcript variant X2, mRNA | 0.034844927 | 0.398787 | 0.039703125 | 0.379961 | 8.47E-05 | 0.446359 | Cellular events-associated |
| MSTRG.8549.4 | XM_013313437 | PREDICTED: Papilio xuthus UDP-glucuronosyltransferase 2A3-like (LOC106118703), mRNA | 3.91E-05 | 0.67598 | 5.23E-07 | 2.164552 | 2.24E-62 | 67.231781 | Cellular events-associated |
| MSTRG.8972.2 | XM_013282955 | PREDICTED: Papilio polytes juvenile hormone epoxide hydrolase-like (LOC106103250), mRNA | 0.000488104 | 0.346014 | 1.15E-05 | 0.534098 | 1.46E-07 | 3.568838 | Cellular events-associated |
| MSTRG.9268.1 | XM_013282544 | PREDICTED: Papilio polytes inactive pancreatic lipase-related protein 1-like (LOC106102924), mRNA | 1.58E-06 | 47.418743 | 1.64E-06 | 4.60672 | 1.66E-05 | 2.406173 | Cellular events-associated |
| MSTRG.9546.6 | XM_013281612 | PREDICTED: Papilio polytes tafazzin homolog (LOC106102199), mRNA | 0.018944149 | 0.600374 | 6.20E-16 | 0.9412 | 0.000735123 | 0.608251 | Cellular events-associated |
| MSTRG.9695.2 | XM_013279222 | PREDICTED: Papilio polytes ubiquitin-protein ligase E3C (LOC106100394), mRNA | 6.45E-07 | 1.288144 | 1.04E-06 | 0.264561 | 5.21E-15 | 0.712137 | Cellular events-associated |
| PmemnonGene0003795.mrna1 | XM_013279734 | PREDICTED: Papilio polytes probable elongation factor 1-delta (LOC106100731), transcript variant X2, mRNA | 8.17E-08 | 38.052101 | 2.52E-21 | 17.745222 | 3.01E-32 | 30.855316 | Cellular events-associated |
| PmemnonGene0004061.mrna1 | XM_014502528 | PREDICTED: Papilio machaon solute carrier organic anion transporter family member 4A1-like (LOC106710467), mRNA | 2.25E-10 | 2.138026 | 4.50E-06 | 1.254498 | 1.81E-05 | 0.59437 | Cellular events-associated |
| PmemnonGene0004994.mrna1 | XM_013278915 | PREDICTED: Papilio polytes dipeptidyl peptidase 3 (LOC106100162), transcript variant X2, mRNA | 3.38E-52 | 2.770487 | 1.08E-14 | 3.687592 | 1.11E-07 | 2.628449 | Cellular events-associated |
| PmemnonGene0005113.mrna1 | XM_013278813 | PREDICTED: Papilio polytes elongation of very long chain fatty acids protein 7-like (LOC106100077), mRNA | 2.58E-06 | 2.999784 | 1.50E-16 | 0.95989 | 1.53E-12 | 16.458216 | Cellular events-associated |
| PmemnonGene0005539.mrna1 | XM_013278653 | PREDICTED: Papilio polytes protein yippee-like CG15309 (LOC106099958), transcript variant X3, mRNA | 9.25E-26 | 3.260163 | 8.81E-08 | 0.274515 | 2.41E-09 | 0.178785 | Cellular events-associated |
| PmemnonGene0006623.mrna1 | XM_013287185 | PREDICTED: Papilio polytes glycine-rich protein DOT1-like (LOC106106584), mRNA | 0.002480876 | 1.683437 | 0.039703125 | 0.469653 | 5.95E-06 | 8.204401 | Cellular events-associated |
| PmemnonGene0008303.mrna1 | XM_013278787 | PREDICTED: Papilio polytes longitudinals lacking protein, isoforms H/M/V-like (LOC106100050), transcript variant X3, mRNA | 6.39E-33 | 3.196245 | 7.11E-40 | 1.155066 | 1.56E-46 | 2.147843 | Cellular events-associated |
| PmemnonGene0010236.mrna1 | XM_013285314 | PREDICTED: Papilio polytes aladin-like (LOC106105089), mRNA | 0.003938964 | 0.550008 | 0.022219062 | 0.237044 | 4.27E-10 | 0.272352 | Cellular events-associated |
| PmemnonGene0002468.mrna1 | AK402281 | Papilio polytes mRNA for cuticular protein PpolCPG14, complete cds, sequence id: Pp-0034 | 1.59E-13 | 10.374036 | 5.70E-11 | 9.903081 | 1.86E-22 | 108.754219 | Cuticular protein |
| PmemnonGene0008556.mrna1 | AK402473 | Papilio polytes mRNA for cuticular protein PpolCPH4A, complete cds, sequence id: Pp-0299 | 1.44E-50 | 594.014282 | 4.29E-19 | 611.811584 | 2.84E-21 | 328.103088 | Cuticular protein |
| PmemnonGene0008557.mrna1 | XM_013281214 | PREDICTED: Papilio polytes pupal cuticle protein PCP52-like (LOC106101864), mRNA | 1.01E-83 | 861.065979 | 1.39E-124 | 946.433105 | 5.86E-24 | 25.847094 | Cuticular protein |
| PmemnonGene0011670.mrna1 | XM_013280577 | PREDICTED: Papilio polytes larval/pupal rigid cuticle protein 66-like (LOC106101375), transcript variant X2, mRNA | 9.39E-35 | 33.526108 | 2.71E-12 | 133.707413 | 1.07E-25 | 108.477745 | Cuticular protein |
| PmemnonGene0011671.mrna1 | XM_013280576 | PREDICTED: Papilio polytes larval/pupal rigid cuticle protein 66-like (LOC106101375), transcript variant X1, mRNA | 1.74E-36 | 33.20509 | 2.33E-14 | 150.683716 | 1.63E-27 | 165.167389 | Cuticular protein |
| PmemnonGene0011676.mrna1 | NM_001311447 | Papilio polytes pupal cuticle protein Edg-84A-like (LOC106101304), mRNA | 7.61E-10 | 5208.787598 | 3.21E-12 | 3892.594238 | 1.23E-06 | 736.309937 | Cuticular protein |
| MSTRG.1168.1 | NM_001311577 | Papilio polytes insecticyanin-B-like (LOC106101117), mRNA | 1.63E-37 | 6224.285645 | 9.15E-36 | 21751.55664 | 2.74E-22 | 46921.14844 | Pigment binding |
| MSTRG.1168.4 | NM_001311545 | Papilio polytes bilin-binding protein-like (LOC106101099), mRNA | 3.16E-35 | 1672.952759 | 3.12E-54 | 10318.94141 | 1.79E-24 | 9616.539062 | Pigment binding |
| MSTRG.1174.1 | NM_001311600 | Papilio polytes bilin-binding protein-like (LOC106101135), mRNA | 1.86E-106 | 281.600281 | 4.20E-93 | 1474.666382 | 4.30E-23 | 164.744446 | Pigment binding |
| PmemnonGene0012312.mrna1 | NM_001311545 | Papilio polytes bilin-binding protein-like (LOC106101099), mRNA | 1.00E-47 | 9093.069336 | 7.68E-09 | 7773.87793 | 4.18E-27 | 28020.58789 | Pigment binding |
| PmemnonGene0012313.mrna1 | AK405661 | Papilio polytes mRNA, putative 3'UTR of hypothetical protein, sequence id: Pp-1323, expressed in epidermis | 4.51E-72 | 451.837494 | 9.13E-67 | 2041.56604 | 3.21E-16 | 170.259644 | Pigment binding |

### Continued Table S2. Green color-associated DEGs.

| Transcripts ID | Assession No. | Description | PValue | FPKM (3rd_P) | PValue | FPKM (4th_P) | PValue | FPKM (5th_A) |
|---|---|---|---|---|---|---|---|---|
| MSTRG.11488.2 | XM_013310070 | PREDICTED: Papilio xuthus protein toll-like (LOC106116294), transcript variant X7, mRNA | 5.56E-14 | 3.736879 | 6.52E-06 | 1.306447 | 0.001226077 | 0.597096 |
| MSTRG.1445.1 | XM_013279332 | PREDICTED: Papilio polytes FGFR1 oncogene partner 2 homolog (LOC106100469), mRNA | 8.71E-11 | 1.234689 | 7.94E-12 | 4.201031 | 2.54E-06 | 2.430666 |
| MSTRG.2313.6 | XM_013287351 | PREDICTED: Papilio polytes NF-kappa-B inhibitor cactus-like (LOC106106715), mRNA | 3.93E-114 | 26.96184 | 3.33E-33 | 45.004093 | 3.09E-109 | 20.129786 |
| MSTRG.2809.5 | XM_013281785 | PREDICTED: Papilio polytes ejaculatory bulb-specific protein 3-like (LOC106102350), mRNA | 0.012496713 | 1.157161 | 1.63E-06 | 2.862453 | 1.26E-08 | 14.927616 |
| MSTRG.2844.3 | XM_013281684 | PREDICTED: Papilio polytes ejaculatory bulb-specific protein 3-like (LOC106102259), mRNA | 1.33E-12 | 1.70846 | 1.26E-20 | 6.815357 | 6.77E-70 | 26.91004 |
| PmemnonGene0000429.mrna1 | XM_013280806 | PREDICTED: Papilio polytes protein sprint (LOC106101556), mRNA | 0.006606612 | 0.191748 | 1.97E-12 | 0.517949 | 0.010399363 | 0.104254 |
| PmemnonGene0000673.mrna1 | XM_013285211 | PREDICTED: Papilio polytes sorting nexin lst-4 (LOC106105012), transcript variant X2, mRNA | 4.71E-12 | 1.48054 | 3.24E-22 | 1.497464 | 0.000262323 | 0.855064 |
| PmemnonGene0004235.mrna1 | XM_013283980 | PREDICTED: Papilio polytes neurogenic locus protein delta-like (LOC106104042), mRNA | 5.43E-10 | 0.990149 | 1.14E-07 | 0.5801 | 1.37E-15 | 13.068659 |
| PmemnonGene0004969.mrna1 | XM_013285612 | PREDICTED: Papilio polytes PDZ and LIM domain protein Zasp-like (LOC106105322), mRNA | 1.59E-06 | 20.174774 | 1.84E-42 | 8.64592 | 1.69E-06 | 13.731862 |
| PmemnonGene0006583.mrna1 | XM_013291027 | PREDICTED: Papilio polytes discoidin domain-containing receptor 2-like (LOC106109507), transcript variant X2, mRNA | 7.36E-08 | 20.876806 | 3.46E-44 | 10.308157 | 5.29E-09 | 51.756653 |
| MSTRG.4559.3 | XM_013280411 | PREDICTED: Papilio polytes transcription elongation regulator 1-like (LOC106101251), mRNA | 1.14E-24 | 0.932547 | 4.24E-34 | 4.444982 | 3.95E-38 | 1.982885 |
| MSTRG.5708.1 | XM_013278075 | PREDICTED: Papilio polytes heat shock transcription factor-like (LOC106099512), mRNA | 1.09E-08 | 4.414275 | 6.18E-09 | 1.860237 | 1.86E-05 | 2.157268 |
| MSTRG.6311.7 | XM_013292623 | PREDICTED: Papilio polytes protein split ends-like (LOC106110726), mRNA | 7.56E-31 | 1.155151 | 0.000447975 | 0.083772 | 0.002426099 | 0.094819 |
| MSTRG.10513.5 | XM_014503471 | PREDICTED: Papilio machaon uncharacterized LOC106711208 (LOC106711208), mRNA | 0.008680466 | 0.397742 | 1.16E-06 | 2.877703 | 7.69E-06 | 1.149921 |
| MSTRG.10893.2 | XM_013280993 | PREDICTED: Papilio polytes uncharacterized LOC106101673 (LOC106101673), transcript variant X2, mRNA | 1.59E-05 | 0.614408 | 1.44E-07 | 0.778773 | 1.89E-05 | 0.464954 |
| MSTRG.1169.2 | XM_013290336 | PREDICTED: Papilio polytes uncharacterized LOC106108964 (LOC106108964), mRNA | 1.81E-09 | 1.087341 | 2.23E-29 | 8.343893 | 3.72E-36 | 52.791115 |
| MSTRG.11723.1 | XM_013292517 | PREDICTED: Papilio polytes uncharacterized LOC106110630 (LOC106110630), transcript variant X2, mRNA | 6.11E-05 | 1.218963 | 0.039703125 | 0.474344 | 0.000556462 | 0.524942 |
| MSTRG.12280.4 | XM_013314700 | PREDICTED: Papilio xuthus uncharacterized LOC106119644 (LOC106119644), mRNA | 1.31E-08 | 3.536931 | 5.57E-06 | 2.139514 | 4.78E-12 | 1.478091 |
| MSTRG.3386.1 | XR_001228180 | PREDICTED: Papilio xuthus uncharacterized LOC106124035 (LOC106124035), ncRNA | 0.03847433 | 0.297234 | 3.83E-23 | 4.801345 | 3.46E-35 | 11.612783 |
| MSTRG.4056.1 | FR989965 | Anthocharis cardamines genome assembly, chromosome: 15 | 0.013197567 | 0.827924 | 1.12E-22 | 11.003154 | 2.92E-33 | 50.425564 |
| MSTRG.4413.3 | XM_013277956 | PREDICTED: Papilio polytes uncharacterized LOC106099404 (LOC106099404), mRNA | 2.73E-09 | 0.170129 | 3.64E-40 | 3.401441 | 6.51E-08 | 1.371988 |
| MSTRG.4817.3 | AK405661 | Papilio polytes mRNA, putative 3'UTR of hypothetical protein, sequence id: Pp-1323, expressed in epidermis | 3.91E-11 | 5.999988 | 7.37E-40 | 3.852819 | 6.91E-20 | 2.922457 |
| MSTRG.5122.2 | XM_014514345 | PREDICTED: Papilio machaon uncharacterized LOC106719873 (LOC106719873), mRNA | 1.18E-06 | 12.643449 | 1.13E-110 | 48.587181 | 1.75E-08 | 63.093906 |
| MSTRG.7916.1 | AK405661 | Papilio polytes mRNA, putative 3'UTR of hypothetical protein, sequence id: Pp-1323, expressed in epidermis | 3.35E-17 | 33.920372 | 0.000627583 | 2.589772 | 8.01E-19 | 8.065794 |
| MSTRG.7999.12 | XM_013290560 | PREDICTED: Papilio polytes uncharacterized LOC106109156 (LOC106109156), mRNA | 6.55E-52 | 4.108368 | 2.63E-09 | 4.340475 | 5.46E-28 | 3.795538 |
| MSTRG.9545.2 | XM_013307077 | PREDICTED: Papilio xuthus enhancer of mRNA-decapping protein 3 (LOC106114017), mRNA | 7.85E-07 | 0.518476 | 1.18E-07 | 1.4503 | 2.50E-16 | 0.865322 |
| PmemnonGene0000577.mrna1 | FR997745 | Ochropleura plecta genome assembly, chromosome: 23 | 1.72E-18 | 336.731689 | 7.68E-15 | 232.074982 | 6.10E-26 | 82.27729 |
| PmemnonGene0001622.mrna1 | XM_013316965 | PREDICTED: Papilio xuthus methyltransferase-like protein 9 (LOC106121340), transcript variant X1, mRNA | 0.000206754 | 0.872263 | 0.010578883 | 0.79658 | 2.10E-07 | 1.672393 |
| PmemnonGene0001661.mrna1 | XM_013284613 | PREDICTED: Papilio polytes uncharacterized LOC106104536 (LOC106104536), mRNA | 1.62E-34 | 16.988594 | 1.25E-06 | 21.289095 | 1.08E-13 | 7.792877 |
| PmemnonGene0001662.mrna1 | XM_013284613 | PREDICTED: Papilio polytes uncharacterized LOC106104536 (LOC106104536), mRNA | 8.01E-34 | 31.713257 | 1.09E-09 | 34.143429 | 8.92E-19 | 13.949715 |
| PmemnonGene0002008.mrna1 | XM_013283492 | PREDICTED: Papilio polytes apolipoprotein D-like (LOC106103666), mRNA | 1.35E-08 | 3.092177 | 0.000538084 | 0.783644 | 0.002421522 | 0.936618 |
| PmemnonGene0007988.mrna1 | XM_013293265 | PREDICTED: Papilio polytes uncharacterized LOC106111240 (LOC106111240), mRNA | 3.17E-09 | 2.562959 | 4.55E-05 | 0.958914 | 0.00057344 | 0.250452 |
| PmemnonGene0011565.mrna1 | XM_013278609 | PREDICTED: Papilio polytes Krueppel homolog 1-like (LOC106099917), mRNA | 1.27E-18 | 6.523467 | 3.68E-37 | 14.461278 | 2.07E-14 | 5.183138 |
| PmemnonGene0012296.mrna1 | XM_013280211 | PREDICTED: Papilio polytes protein kinase C (LOC106101096), transcript variant X4, mRNA | 1.54E-05 | 2.350068 | 0.000398911 | 1.112151 | 3.38E-09 | 0.708404 |

**Table S3. siRNA list.**

| Target gene | siRNA name | | Sequence |
|---|---|---|---|
| *abdominal-A* | *PmeGene0007395_abd-A* | Target (DNA) | CTCTTCAATCGTTCTAACAGTTC |
| | | Antisense strand (RNA) | ACUGUUAGAACGAUUGAAGAG |
| | | Sense strand (RNA) | CUUCAAUCGUUCUAACAGUUC |
| *Abdominal-B* | *PmeGene0007396_Abd-B* | Target (DNA) | TCCTCTTCAACGCATACGTGTCG |
| | | Antisense strand (RNA) | ACACGUAUGCGUUGAAGAGGA |
| | | Sense strand (RNA) | CUCUUCAACGCAUACGUGUCG |
| *clawless* | *PmeGene0005243_cll* (*Pp*) | Target (DNA) | TACTGCTTTAAGCGCTTTACAAA |
| | | Antisense strand (RNA) | UGUAAAGCGCUUAAAGCAGUA |
| | | Sense strand (RNA) | CUGCUUUAAGCGCUUUACAAA |
| *Antennapedia* | *PmeGene7391_Antp* | Target (DNA) | GCGATAGCATGACATACTTCTCC |
| | | Antisense strand (RNA) | AGAAGUAUGUCAUGCUAUCGC |
| | | Sense strand (RNA) | GAUAGCAUGACAUACUUCUCC |
| *Ultrabithorax* | *Pme_Ubx*_A (7392/739) | Target (DNA) | CAGATATCAGACGCTAGAATTAG |
| | | Antisense strand (RNA) | AAUUCUAGCGUCUGAUAUCUG |
| | | Sense strand (RNA) | GAUAUCAGACGCUAGAAUUAG |
| *delta* | *PmeGene0004235_Delta_P.p* | Target (DNA) | ACGATTCATTCGGACATTACACC |
| | | Antisense strand (RNA) | UGUAAUGUCCGAAUGAAUCGU |
| | | Sense strand (RNA) | GAUUCAUUCGGACAUUACACC |
| *Bilin-binding protein 1* | *PmeBBP1* | Target (DNA) | AGGTTATGCTGACATACACATTT |
| | | Antisense strand (RNA) | AUGUGUAUGUCAGCAUAACCU |
| | | Sense strand (RNA) | GUUAUGCUGACAUACACAUUU |
| *Bilin-binding protein 2* | *PmeBBP2* | Target (DNA) | GAGGAGATAACCGAATTCTTAAA |
| | | Antisense strand (RNA) | UAAGAAUUCGGUUAUCUCCUC |
| | | Sense strand (RNA) | GGAGAUAACCGAAUUCUUAAA |

**Continued Table S3. siRNA list.**

| Target gene | siRNA name | | Sequence |
|---|---|---|---|
| *Bilin-binding protein 3* | *Pme_BBP3* | Target (DNA) | AAGTAGTGAACGGCGTCAAATCT |
| | | Antisense strand (RNA) | AUUUGACGCCGUUCACUACUU |
| | | Sense strand (RNA) | GUAGUGAACGGCGUCAAAUCU |
| *Bilin-binding protein 4* | *Pme_BBP4* | Target (DNA) | AACTTTGATTTCAATGCTTATCA |
| | | Antisense strand (RNA) | AUAAGCAUUGAAAUCAAAGUU |
| | | Sense strand (RNA) | CUUUGAUUUCAAUGCUUAUCA |
| *Bilin-binding protein 5* | *Pme_BBP5* | Target (DNA) | AACAGAACTTCGATTTTGCATCT |
| | | Antisense strand (RNA) | AUGCAAAAUCGAAGUUCUGUU |
| | | Sense strand (RNA) | CAGAACUUCGAUUUUGCAUCU |
| *Bilin-binding protein 6* | *Pme_BBP6* | Target (DNA) | CTCTAATCGAGTACTACGTATAT |
| | | Antisense strand (RNA) | AUACGUAGUACUCGAUUAGAG |
| | | Sense strand (RNA) | CUAAUCGAGUACUACGUAUAU |
| *JH-binding protein 4811* | *Pme_JHBP4811* | Target (DNA) | TTGAATCTACTGGCAAATACAAG |
| | | Antisense strand (RNA) | UGUAUUUGCCAGUAGAUUCAA |
| | | Sense strand (RNA) | GAAUCUACUGGCAAAUACAAG |
| *JH-binding protein 4812* | *Pme_JHBP4812* | Target (DNA) | AAGCGTTACTGGAAACTGAATCT |
| | | Antisense strand (RNA) | AUUCAGUUUCCAGUAACGCUU |
| | | Sense strand (RNA) | GCGUUACUGGAAACUGAAUCU |
| *JH-binding protein 4813* | *Pme_JHBP4813* | Target (DNA) | CTCTACTATGTCCGTAACTTTAG |
| | | Antisense strand (RNA) | AAAGUUACGGACAUAGUAGAG |
| | | Sense strand (RNA) | CUACUAUGUCCGUAACUUUAG |
| *JH-binding protein 4814* | *Pme_JHBP4814* | Target (DNA) | GTGATGAATCTGCTATGAGATTC |
| | | Antisense strand (RNA) | AUCUCAUAGCAGAUUCAUCAC |
| | | Sense strand (RNA) | GAUGAAUCUGCUAUGAGAUUC |
| *JH-binding protein 4815* | *Pme_JHBP4815* | Target (DNA) | CTCCTAAGTTTCGAATACTCAAG |
| | | Antisense strand (RNA) | UGAGUAUUCGAAACUUAGGAG |
| | | Sense strand (RNA) | CCUAAGUUUCGAAUACUCAAG |
| *JH-binding protein 4816* | *Pme_JHBP4816* | Target (DNA) | TACCTTCAACGATCTAAACATAT |
| | | Antisense strand (RNA) | AUGUUUAGAUCGUUGAAGGUA |
| | | Sense strand (RNA) | CCUUCAACGAUCUAAACAUAU |
| *JH-binding protein 2505* | *Pme_JHBP2505* | Target (DNA) | AACCGCTAATCATGTTCTTAAAT |
| | | Antisense strand (RNA) | UUAAGAACAUGAUUAGCGGUU |
| | | Sense strand (RNA) | CCGCUAAUCAUGUUCUUAAAU |
| *JH-binding protein 2506* | *Pme_JHBP2506* | Target (DNA) | AAGGAAAACTCCTGATTAAGTTG |
| | | Antisense strand (RNA) | ACUUAAUCAGGAGUUUUCCUU |
| | | Sense strand (RNA) | GGAAAACUCCUGAUUAAGUUG |

**Table S4. Primer list.** (Continued on next page)

| Target gene | Primer name | Sequence |
|---|---|---|
| *abdominal-A* | *P. me abd-A* F | CAGGATGTACCCCTACGTGTC |
| | *P. me abd-A* R | GTCAGGGCGTAGTTCATCATGG |
| *Abdominal-B* | *P. me Abd-B* F | CCCCACATACTACAATCTGCCA |
| | *P. me Abd-B* R | GGCTGAGAGAAACCCTGATGAC |
| *clawless* | *P. me cll* F | GGCTGAAGCGCTAAGTAAGGG |
| | *P. me cll* R | TAAAGCAGTATTCGCTGGAGGC |
| *ribosomal protein L3* | *P. me rpL3* F | CTTGAGAAGCCTATCCCAGTGG |
| | *P. me rpL3* R | GAGCTTTTTAGTGTGCCAACGG |
| *Bilin-binding protein 1* | *Pme_BBP1*_F | TACGGTGGGACATGGTACGA |
| | *Pme_BBP1*_R | TGAATTCCTCGCTTTGCCCT |
| *Bilin-binding protein 2* | *Pme_BBP2*_F | AATCGCCAAATTCCCAAACCC |
| | *Pme_BBP2*_R | CTTGCGCCATCAACTACGTG |
| *Bilin-binding protein 3* | *Pme_BBP3*_F | GGTAAATGCTCCACTGCGGA |
| | *Pme_BBP3*_R | TCCAGGTCCCACAAGAGTCA |
| *Bilin-binding protein 4* | *Pme_BBP4*_F | TGAAGCCGATGGAGAACTTTGA |
| | *Pme_BBP4*_R | GTGTTTCTAACTTTGCCGCTGT |
| *Bilin-binding protein 5* | *Pme_BBP5*_F | CCTGTCCGTCTGTCAGACCT |
| | *Pme_BBP5*_R | CAATCGTGCACTTTCCCCTG |
| *Bilin-binding protein 6* | *Pme_BBP6*_F | TTGTTCGTGTTTGCTCTGATCG |
| | *Pme_BBP6*_R | ATGTTCCTTGGTAAGCGGAGAA |

**Continued Table S4. Primer list.**

| Target gene | Primer name | Sequence |
|---|---|---|
| *JH-binding protein 4811* | *Pme_JHBP4811*_F | CCGAATATGAAGTTAACCGGCG |
| | *Pme_JHBP4811*_R | CCAGTAGATTCAAACGGACAGTC |
| *JH-binding protein 4812* | *Pme_JHBP4812*_F | TTCTTTAGCCCGTCGACTGG |
| | *Pme_JHBP4812*_R | TGGTACACGTTGGTAGAATGCA |
| *JH-binding protein 4813* | *Pme_JHBP4813*_F | GCCCTCTCTAGAATCGGTTATGG |
| | *Pme_JHBP4813*_R | AGTTACGGACATAGTAGAGCGTG |
| *JH-binding protein 4814* | *Pme_JHBP4814*_F | TCCCACCGACAGATCCGAT |
| | *Pme_JHBP4814*_R | ATTCAACGACCCTGCATCTCC |
| *JH-binding protein 4815* | *Pme_JHBP4815*_F | TGTCGTCAACAACTCCAGCA |
| | *Pme_JHBP4815*_R | ATTCCCTCGGCGATGTATGG |
| *JH-binding protein 4816* | *Pme_JHBP4816*_F | ATAACATCCAGTGCTACGGC |
| | *Pme_JHBP4816*_R | TTTCATGTCACCCTGGTCGG |
| *JH-binding protein 2505* | *Pme_JHBP2505*_F | AGTCGCAGGAGAACTCGTTG |
| | *Pme_JHBP2505*_R | TGACCGTCGCCATCTAAAATCT |
| *JH-binding protein 2506* | *Pme_JHBP2506*_F | CAGTGCCCGTCAACAAGATATG |
| | *Pme_JHBP2506*_R | TCAAATTGGGTGAACTTGCGTC |

**Table S5. Electroporation conditions for the *in vivo* RNAi.**

| Poring Pulse | | Transfer Pulse | |
|---|---|---|---|
| Voltage | 20 V | Voltage | 15 V |
| Pulse Length | 5 msec | Pulse Length | 90 msec |
| Pulse Interval | 20 msec | Pulse Interval | 50 msec |
| Number of Pulse | 2 | Number of Pulse | 20 |
